# *Engrailed-1* Promotes Pancreatic Cancer Metastasis

**DOI:** 10.1101/2023.04.10.536259

**Authors:** Jihao Xu, Jae-Seok Roe, EunJung Lee, Claudia Tonelli, Tim D.D. Somerville, Melissa Yao, Joseph P. Milazzo, Herve Tiriac, Ania M. Kolarzyk, Esak Lee, Jean L. Grem, Audrey J. Lazenby, James A. Grunkemeyer, Michael A. Hollingsworth, Alexander D. Borowsky, Youngkyu Park, Christopher R. Vakoc, David A. Tuveson, Chang-Il Hwang

## Abstract

*Engrailed-1* (*EN1*) is a critical homeodomain transcription factor (TF) required for neuronal survival, and EN1 expression has been shown to promote aggressive forms of triple negative breast cancer. Here, we report that *EN1* is aberrantly expressed in a subset of pancreatic ductal adenocarcinoma (PDA) patients with poor outcomes. EN1 predominantly repressed its target genes through direct binding to gene enhancers and promoters, implicating a role in the acquisition of mesenchymal cell properties. Gain- and loss-of-function experiments demonstrated that *EN1* promoted PDA transformation and metastasis *in vitro* and *in vivo*. Our findings nominate the targeting of *EN1* and downstream pathways in aggressive PDA.

## Introduction

Pancreatic cancer is the third leading cause of cancer-related deaths in the United States with 12% 5-year relative survival rate, lowest among all common cancers (1). Pancreatic ductal adenocarcinoma (PDA) is the most common and challenging form of pancreatic cancer due to its highly metastatic nature, lack of screening, and resistance to current chemotherapeutic regimens. In PDA, carcinogenesis begins with a gain-of-function mutation of *KRAS* in the pancreatic ductal epithelial cells, leading to the formation of pancreatic intraepithelial neoplasia (PanIN) lesions. The activating mutation in *KRAS* then cooperates with the loss-of-function mutations of tumor suppressor genes, including *TP53*, *SMAD4*, and *CKDN2A*, to further promote PDA progression (2). In contrast, recurrent genetic alterations driving PDA metastasis remains elusive. Instead, metastatic lesions harbor a similar pattern of driver mutations as seen throughout the primary PDA (3), suggesting that PDA metastasis may be driven by nongenetic alterations, such as fluctuations in signal transduction and transcriptional programs. However, the molecular mechanisms driving such fluctuations are understudied; therefore, there is an urgent need to understand the driving force behind PDA progression in order to develop new therapeutic strategies to counter disease progression and improve patients’ survival.

Aberrant expressions of transcription factors (TFs) and the subsequent alterations in epigenetic landscapes may be responsible for the fluctuations of transcriptional programs during cancer progression (4). For example, through modulating enhancer activities, TF TP63 is capable to activate the transcriptional programs of squamous PDA subtype, leading to an aggressive cancer phenotype (5). Until recently, studying the dynamic changes of transcriptional programs as the cancer progresses has been difficult due to lack of *in vitro* PDA progression models for different stages of PDA. To address this, we previously established an *in vitro* organoid model derived from *Kras^+/LSL-G12D^; Trp53^+/LSL-R172H^; Pdx1-Cre* (KPC) mouse (6). This model allowed a direct comparison of the tumor (mT)- and paired metastasis (mM)-derived organoids from the KPC mouse and showed that TF FOXA1 is capable to activate the transcriptional programs of endoderm lineage through enhancer reprogramming, promoting PDA metastasis (7). Likewise, it is possible that other TFs are also aberrantly regulated in PDA progression and confer aggressive characters through such epigenetic reprogramming, which warrants a further investigation.

During the development, the required cellular processes (e.g., differentiation and death) are tightly regulated by interactions between epigenomes and TF-mediated lineage-specific gene programs (8). Interestingly, the genes involved in neurodevelopmental programs, such as axon guidance pathways, are frequently altered in many cancers, including PDA, leading to the disease progression (9). It is therefore probable that cancer cells hijack TFs that govern these developmental pathways to confer a survival benefit. Homeobox TFs are evolutionarily conserved master regulators that are essential for embryonic development. Among the homeobox TFs, Engrailed-1 (EN1) is essential in the development of central nervous system and implicated in the control of cell differentiation, growth, survival, and axon guidance at the cellular level (10, 11). In addition, a number of studies have reported aberrantly expressed EN1 and the association with poor prognosis in human malignancies, including glioblastoma, salivary gland adenoid cystic carcinoma, and breast-related cancers (12–18). However, the detailed molecular mechanisms by which EN1 promotes PDA progression remain unknown.

In this study, we showed that EN1 promotes metastatic properties in PDA, through direct bindings to promoter and enhancer of the genes involved in several cellular pathways, including cell death and mitogen-activated protein kinase (MAPK) pathways. As a result, aberrant expression of EN1 accelerates PDA progression *in vivo*. Therefore, targeting EN1 and its downstream pathways can be effective therapeutic strategies for EN1-high PDA patients.

## Results

### EN1 expression is associated with PDA progression and patient poor prognosis

To identify pro-survival factors contributing to PDA progression, we first developed an organoid survival assay where single cell-dissociated pancreatic organoids were grown in DMEM supplemented with 10% FBS without additional growth factors and Wnt ligands (hereafter referred to as the reduced media). In the reduced media, only mM organoid-derived cells survived, formed organoids, and could be passaged continuously, while mT organoid-derived cells failed to grow (Fig. 1A and S1A-B). Consistent with the known functions of wild-type p53 in cell death (19), the inactivation of p53 in mT organoids resulted in increased organoid forming in the organoid survival assay, and promoted PDA progression *in vivo* (Fig S1C-H), suggesting that organoid survival phenotype can be served as a translatable readout for the *in vivo* context. Through transcriptome and epigenome profiling on the paired mT and mM organoids, we previously identified that aberrant expression of several developmental TFs led to enhancer reprogramming and endowed aggressive characteristics seen in PDA metastasis (7, 20). The TFs, including *Batf2*, *Foxa1*, *Gata5*, *Prrx2*, *Pax9*, *Trerf1*, and *En1*, were highly expressed in mM organoids compared to their paired mT organoids (Fig. 1B and S1I). To identify functionally important TF(s), we performed the organoid survival assay with mT organoids and the 7 TFs. The retroviral introduction of the 7 TFs enabled mT organoids to survive and propagate in the reduced media (Fig. 1C). The withdrawal of an individual TF revealed that mT organoids failed to survive and form organoids without EN1 (Fig. 1C and S1J). Likewise, the introduction of *En1* cDNA in mT organoids increased organoid survival, suggesting that EN1 is a critical pro-survival factor in PDA (Figure 1D and S1K). In concordance with our previous finding (7), when FOXA1 was removed from the 7 TF combination, the organoid survival was also impeded, although to a lesser degree than the EN1 withdrawal. To determine if advanced stage of PDA expressed EN1 proteins, we performed EN1 IHC using the KPC mice tissue sections and the orthotopic transplantation sections of the mT organoids. Indeed, we confirmed that EN1 expression was elevated in the late stage of PDA (Fig. 1E-F and S1L). We then analyzed the publicly available RNA-seq datasets of human PDA patients (21, 22). Consistent with murine models of PDA, we found that *EN1* expression was elevated in the advanced stage of PDA (Fig. 1G and S1M). In addition, analyses of the transcriptomic profile of PDA PDOs and scRNA-seq dataset (23, 24) revealed that a subset of PDA patients showed EN1 expression (Figure 1H and S1N). Moreover, we found that the increased expression of *EN1* was associated with poor prognosis in the TCGA dataset (Fig. 1I). Gene Set Enrichment Analysis (GSEA) of the publicly available expression datasets (21, 24–27) further revealed that *EN1* expression was closely associated with the molecular signatures implicated in aggressiveness of PDA, including epithelial-mesenchymal transition (EMT) and squamous/basal-like molecular subtype (Fig. 1J-M and S1O). Therefore, EN1 is tightly associated with the aggressive features of PDA and aberrant expression of EN1 could provide pro-survival cues and contribute to aggressiveness of PDA cells.

**Figure 1.**
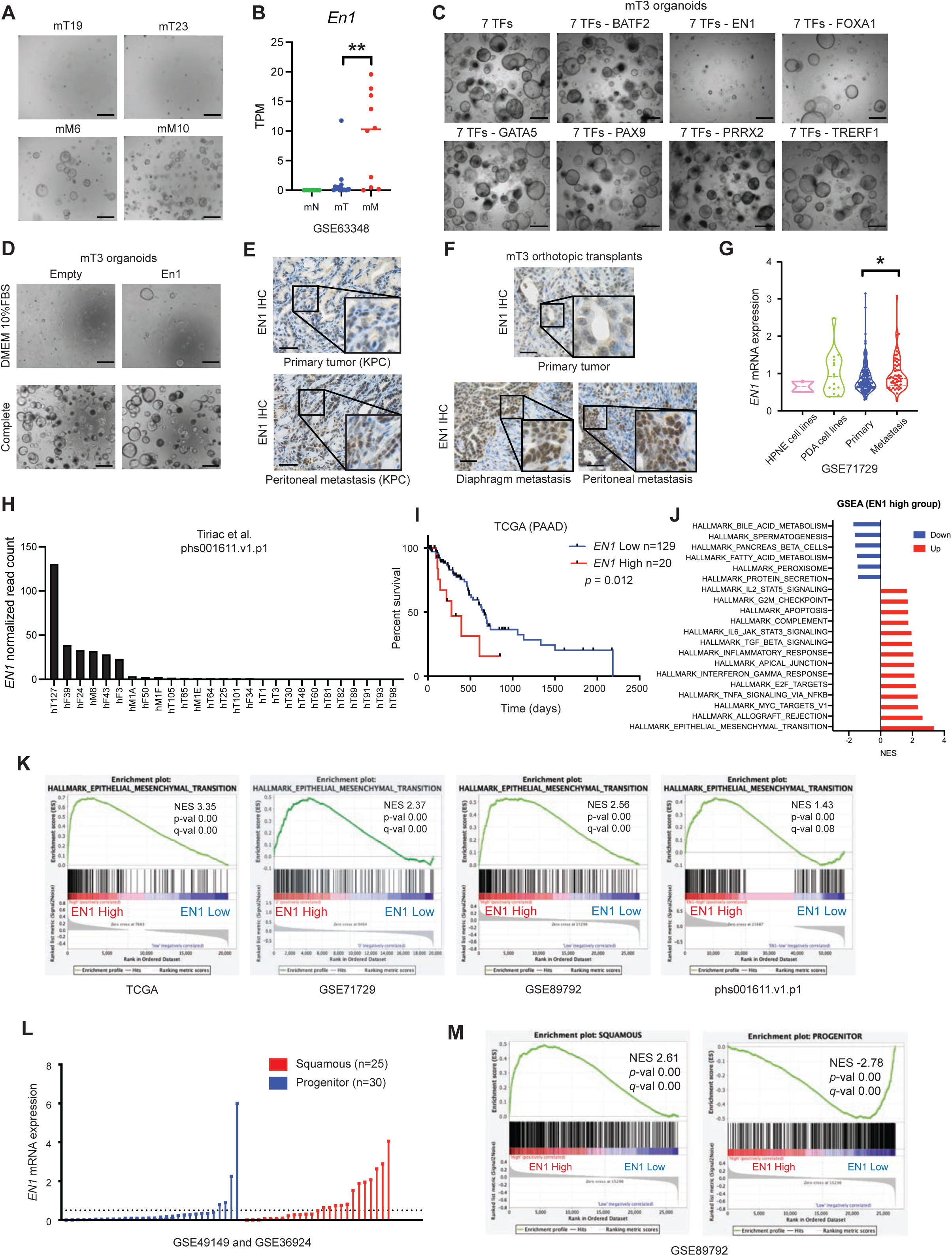
EN1 expression is associated with PDA progression and patient poor prognosis. (A) Organoid survival assay of KPC tumor (mT)- and metastasis (mM)-derived organoids. Organoids were dissociated into single cells, plated and grown in 10% FBS DMEM (reduced media) for 5 days. Scale bars, 1 mm. (B) RNA-seq based *En1* mRNA expression in organoids from Oni et al. (GSE66348). Each dot represents an organoid line. (C) 7 TFs (BATF, EN1, FOXA1, GATA5, PAX9, PRRX2, TRERF1) were introduced in mT3 organoids and subjected to organoid survival assay. Each TF was withdrawn from the 7 TFs combination. Scale bars, 1 mm. (D) mT3 organoids with En1 cDNA were subjected to organoid survival assay either in the reduced media or in the complete media for 5 days. (E-F) EN1 IHC of the indicated tissue sections, including primary tumor and peritoneal metastatic lesions from KPC mice (E) and mT3 organoids orthotopic injection models (F). Scale bars, 100 μm. (G) Microarray based *EN1* mRNA expression in cell lines and human PDA tissues from Moffitt et al. (GSE71729). (H) *EN1* mRNA normalized read count from PDA patient-derived organoids (PDOs) in Tiriac et al. (phs001611.v1.p1). (I) EN1 is associated with patient poor prognosis. Pancreatic cancer patients (TCGA-PAAD) were stratified based on *EN1* expression (*EN1*-high n=20 vs. -low n=129). *p-*value was determined by logrank (Mantel-Cox) test. (J) Top 20 significantly enriched hallmarks of GSEA in *EN1*-high vs. -low patients from TCGA-PAAD. Normalized enrichment score (NES) is shown. (K) GSEA of epithelial-to-mesenchymal transition signatures in *EN1*-high vs. -low pancreatic cancer patients or cell lines from TCGA-PAAD, Monffitt et al. (GSE71729), Bian et al. (GSE89792), and Tiriac et al. (phs001611.v1.p1) NES, *p*-value, and FDR *q*-value were determined by GSEA. (L) Normalized *EN1* gene counts in progenitor and squamous PDA subtypes from Bailey et al. (GSE49149 and GSE36924). A dotted line indicates a cutoff to determine EN1 high vs. low to perform Fisher’s exact test (*p*-val < 0.05). The cutoff was determined by a median value of EN1 expression in the squamous subtype. (M) GSEA of squamous (left) and progenitor (right) signatures in *EN1*-high vs. -low pancreatic cancer patients from Bian et al. (GSE89792). NES, *p*-value, and FDR *q*-value were determined by GSEA. Unless otherwise indicated, *p*-values were determined by student’s *t* test (two-tail) and *, **, ***, **** indicated p-val < 0.05, < 0.01, <0.001, <0.0001, respectively.

### EN1 promotes aggressive characteristics in PDA cells

Given the association between EN1 expression and gene signatures of EMT and squamous subtype, we first hypothesized that EN1 could foster aggressive characteristics of PDA. To this end, we retrovirally introduced *En1* cDNA into murine KPC (mT-2D) cell lines (Fig. S2A), and measured cell proliferation, invasion, migration, and anchorage-independent growth. While EN1 did not change the cell proliferation rate (Fig. S2B), we found EN1 overexpression increased the cell invasion (Fig. 2A), migration (Fig. 2B and S2C), and anchorage-independent growth (Fig. 2C), indicating that EN1 promotes metastatic natures of PDA *in vitro*. To further corroborate the role of EN1 in metastatic transitions, we used a two-channel microfluidic organotypic model (28) to investigate the role of EN1 in the intravasation potential of the cells (Fig. 2D). In the model, the collagen-matrix embedded channel (green) contains mT-2D cells and the other channel (red) contains a biomimetic blood vessel. As a result, EN1-expressing cells invaded the collagen matrix toward the blood vessels at a faster rate compared to the control. Furthermore, tail-vein injections of mT-2D cells revealed that EN1-expressing cells readily colonized in the lung parenchyma (Fig. 2E), suggesting that EN1-mediated pro-survival and pro-migratory/invasive phenotypes conferred the metastatic ability necessary for lung colonization. To test if EN1 plays similar roles in the human PDA cells, we chose CFPAC1 and PaTu 8988s human PDA cell lines that do not express EN1 (Fig. S2D), and retrovirally introduced FLAG-tagged *EN1* cDNA. In accordance with the data from murine mT-2D cells, EN1 overexpression increased clonogenic growth (Fig. 2F and S2F, respectively) and anchorage-independent tumor sphere formation (Fig. 2G and S2G, respectively) in CFPAC1 and PaTu 8988s cells. Taken together, the gain-of-function experiments showed that EN1 fosters aggressive characteristics of PDA, including cell survival, migration, and intravasation.

**Figure 2.**
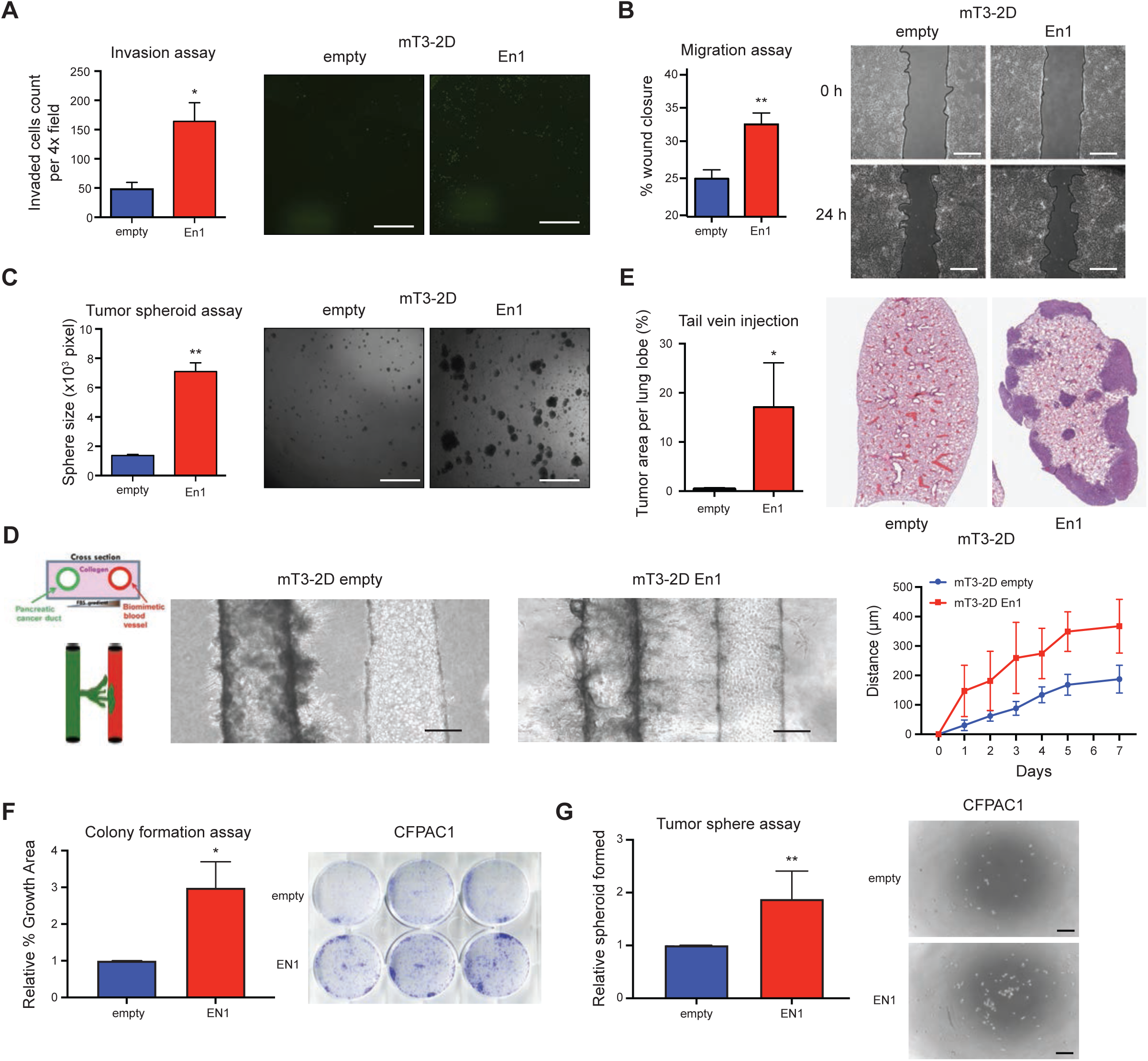
EN1 promotes aggressive characteristics in PDA cells. (A) mT3-2D cells with (En1) and without (empty) *En1* cDNA overexpression were subjected to Boyden-chamber invasion assay for 24 hours, and the cells migrating to across the transwell were stained by SYTO 13 (right) and quantified per 4x image field (left). n=3, mean ± SEM. (B) mT3-2D empty and *En1* cells were subjected to wound-healing assay, and the percentage of wound closure was monitored (right) and quantified (left) at 0- and 24-hour post-scratching. n=3, mean ± SEM. (C) mT3-2D cells with *En1* cDNA were subjected to anchorage-independent tumor spheroid formation assay for 72 hours, and the numbers of spheroids were monitored (right) and quantified (left). n=3, mean ± SEM. (D) mT3-2D empty (n=7) and *En1* (n=4) cells were subjected to organotypic tumor-on-a-chip assay (left) for 7 days, and the distance of the cell migrated toward to endothelial vessel was monitored (middle) and quantified (right) (n=8 per time point, mean ± SD). *p*<0.001; *p*-value were determined by Two-way ANOVA. Scale bar, 200 µm. (E) mT3-2D cells with *En1* cDNA were subject for tail-vein injection (n=5 per group) in C57BL/6 syngeneic mice. After 4 weeks, the animals were sacrificed, and the lung lobes were imaged (right) and quantified (left) for tumor area per lung lobe. n=5, mean ± SD. (F) CFPAC1 empty and *EN1* cells were subject for colony formation assay for 7 days, and the colonies were stained by crystal violet (right) and quantified (left) by percentage growth area. n=9, mean ± SD. (G) CFPAC1 empty and *EN1* cells were subject for anchorage-independent tumor spheroid formation assay for 7 days, and the numbers of spheroids were monitored (right) and quantified (left). n=9, mean ± SD. Scale bars, 350 μm. Unless otherwise indicated, *p*-values were determined by student’s *t* test (two-tail) and * and ** indicate *p*-val < 0.05, and < 0.01, respectively.

### EN1 deficiency attenuates PDA progression

Since EN1 expression contributes to the aggressive natures of PDA cells, we reasoned that EN1-targeting strategy might be therapeutically relevant. To investigate the effects of EN1 depletion in metastatic pancreatic cancer, we lentivirally introduced shRNAs against *En1* either targeting coding sequence (CDS) or 3’-untranslated regions (3’ UTR) into mM3P and mM15 organoids (Fig. S3A). We then subjected the mM organoids to the organoid survival and clonogenic cell growth assays. While EN1 depletion had no effect on cell growth in the complete organoid media (Fig. S3B), we observed that survival of the cells was markedly diminished in the reduced media upon EN1 depletion (Fig. 3A and S3C). Moreover, the ability of clonogenic growth in 2D was also impaired upon *En1* knockdown (Fig. 3B and S3D). To exclude the possibility of shRNA off-target effects, we performed a rescue experiment using retroviral *En1* cDNA. The phenotype of the reduced survival was rescued by *En1* cDNA in the shRNA-3’UTR organoids but not in the shRNA-CDS organoids (Fig. 3A). To confirm the phenotype seen *in vitro*, we used subcutaneous and orthotopic transplantation models of PDA to determine the effects of *En1* knockdown in mM organoids *in vivo*. Previously, we showed mM organoids were highly metastatic compared to the paired mT organoids in orthotopic, tail-vein, and intrasplenic transplantation models (7). Consistent with *in vitro* phenotypes, *En1* knockdown significantly reduced primary tumor burden in both subcutaneous and orthotopic models (Fig. 3C-D). In the orthotopic model, we observed the reduced liver and lung metastases (Fig. 3E-F and S3E-G), suggesting EN1 possibly enhances metastatic potentials of PDA. To address the role of EN1 in the human context, we depleted EN1 using two independent *EN1* shRNA constructs in *EN1*- expressing SUIT2 and BxPC3 human PDA cell lines (Fig. S2D and S3H). As expected, EN1 depletion in human PDA cell lines led to the reduced colony formation and anchorage-independent tumor sphere formation (Fig. 3G-H and S3I-J). Taken together, our results from the loss-of-function experiments showed that EN1 expression is required for cell survival and metastatic capabilities *in vitro* and *in vivo*, suggesting that EN1 and EN1-related pathways might be potential therapeutic targets for PDA.

**Figure 3.**
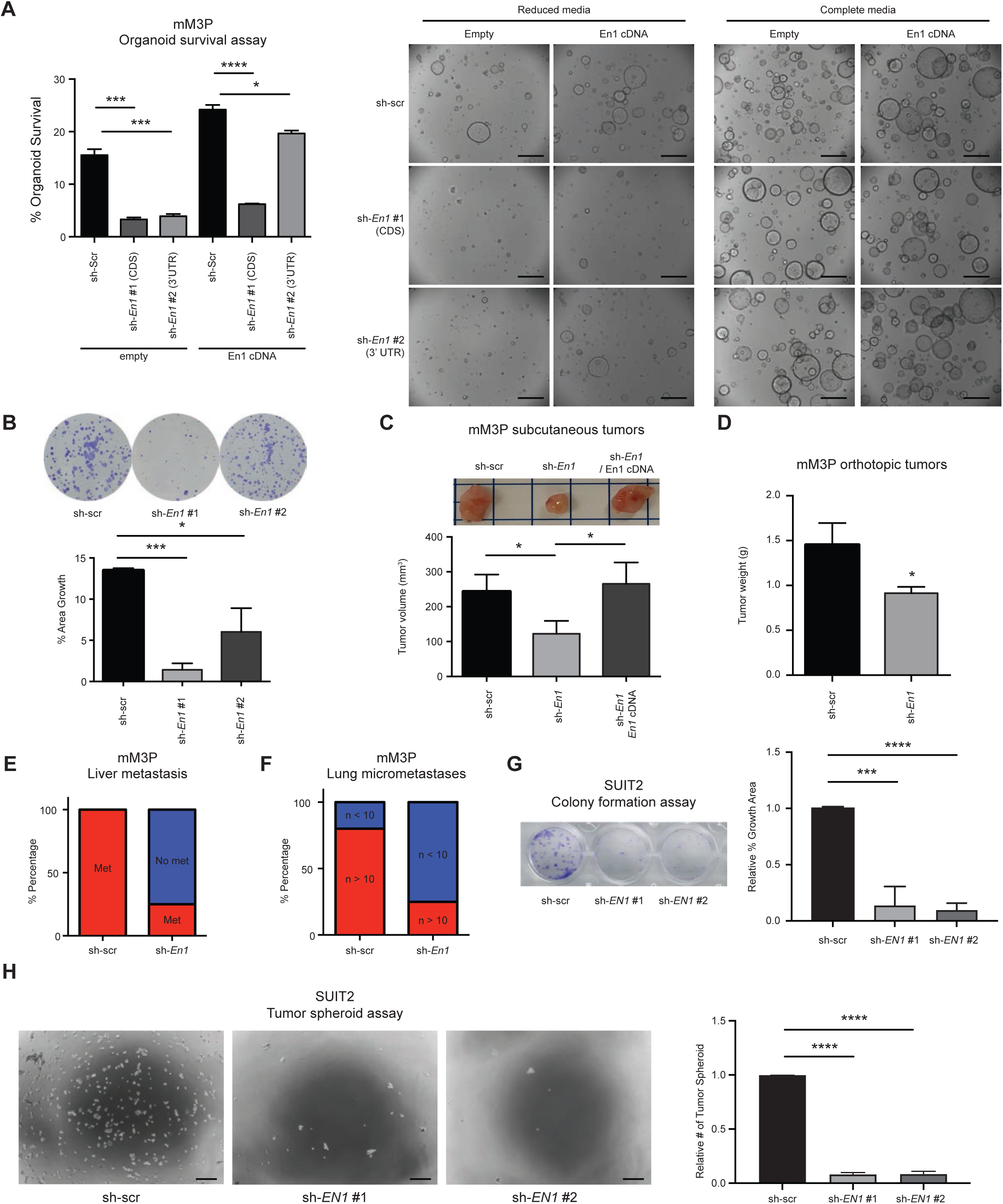
EN1 deficiency attenuates PDA progression. (A) mM3P organoids with scramble (shScr) and two *En1* (sh*En1*) shRNA constructs were subjected to organoid survival assay for 4 days. Depletion of EN1 impaired organoid survival in the reduced media and *En1* cDNA rescued the EN1-depletion phenotype (middle). The complete media was served as control (right). Quantification of organoid survival (left). n=3, mean ± SD. Scale bars, 1mm. (B) shScr and sh*En1* mM3P organoids were subjected to colony formation assay for 7 days, and the colonies were stained by crystal violet (top) and quantified (bottom) by percentage growth area. n=3, mean ± SD. (C) 5×10^5^ cells of dissociated shScr and sh*En1* mM3P organoids were subjected to subcutaneous transplantation in athymic NU/NU mice. The animals were sacrificed at 4-weeks post transplantation. EN1 depletion reduced the primary tumor burden and *En1* cDNA rescued the phenotype. Representative images of the subcutaneous tumors (top) and quantification (bottom) of the tumor volume. n=5 per group, mean ± SD. (D-F) 5×10^5^ cells of dissociated shScr and sh*En1* mM3P organoids were subjected to orthotopic transplantation in athymic NU/NU mice for 7 weeks. The primary tumor weight (D), the number of animals with liver metastases (E), and the number of lungs micrometastasis (n>10) (F) were quantified. The mean ± SD is shown (n=5 for shScr and n=4 for sh*En1*). (G) shScr and sh*EN1* SUIT2 cells were subjected to colony formation assay for 5 days, and the colonies were stained with crystal violet (top) and quantified (bottom) for the percentage growth area. n=3, mean ± SD. (H) shScr and sh*EN1* SUIT2 cells were subjected to anchorage-independent tumor spheroid formation assay for 7 days, and the numbers of spheroids were monitored (left) and quantified (right). n=3, mean ± SD. Scale bars, 350 μm. Unless otherwise indicated, *p*-values were determined by unpaired student’s *t* test (two-tail) and *, **, ***, **** indicate *p*-val < 0.05, < 0.01, <0.001, <0.0001, respectively.

### Identifying genomic targets of EN1 in murine PDA cells

Using gain- and loss-of-function experiments, we thus far demonstrated that EN1 is sufficient and necessary to develop aggressive characteristics in PDA. We reasoned that the underlying mechanisms of how EN1 confers these characteristics are likely dependent on its direct gene targets. Therefore, to dissect the underlying mechanism(s) by which EN1 endows the aggressive characteristics, we attempted to determine genome wide EN1 binding sites and identify direct target genes of EN1. To address this, we first retrovirally introduced FLAG-tagged *En1* cDNA into mT-2D cell lines (Fig. S4A) and performed cleavage under target & release using nuclease followed by sequencing (CUT&RUN-seq) targeting the FLAG epitope in mT4-2D and mT5-2D cell lines. From our bioinformatic analysis of CUT&RUN-seq, we identified 35,256 and 26,582 EN1 peaks in mT4-2D and mT5-2D KPC cells, respectively (Fig. 4A). We then overlapped the two datasets and identified a total of 20,271 common peaks between these two cell lines. Among these peaks, most were located at gene promoters (41.74%) and intergenic/intron regions (54.26%) (Fig. 4B), indicating that EN1 binds at gene promoters and enhancers. HOMER motif analysis of 20,271 common peaks showed the enrichment of known EN1 motif (GSE120957), other homeobox TFs motifs (LHX9 and ISL1) (Fig. S4B), and *de novo* discovery of EN1 motif (Fig. 4C). While the known EN1 motif was enriched in the triple-negative breast cancer (TNBC) cell lines (18), we found minimum overlaps of EN1 peaks between PDA and TNBC cells (Fig. S4C), suggesting EN1 genomic targets could differ depending on tissue or cell types. To understand the functional and biological importance of EN1 genomic targets and peak-associated genes, we performed gene ontology (GO) analysis using Genomic Regions Enrichment of Annotations Tool (GREAT) (Fig. S4C) and the Database for Annotation, Visualization, and Integrated Discovery (DAVID) (Fig. 4D). Both analyses showed the enrichment of apoptotic processes, cytoskeleton organizations, and cell cycle regulations. When the transcriptome profiles of PDA from the publicly available datasets were stratified into *EN1*- high and -low patient groups (24, 29–33), we identified the majority of genes associated with EN1 peaks were down-regulated in *EN1*-high patients (Fig. 4E and S4D), suggesting a predominant transcriptional repressive role of EN1 in PDA cells.

**Figure 4.**
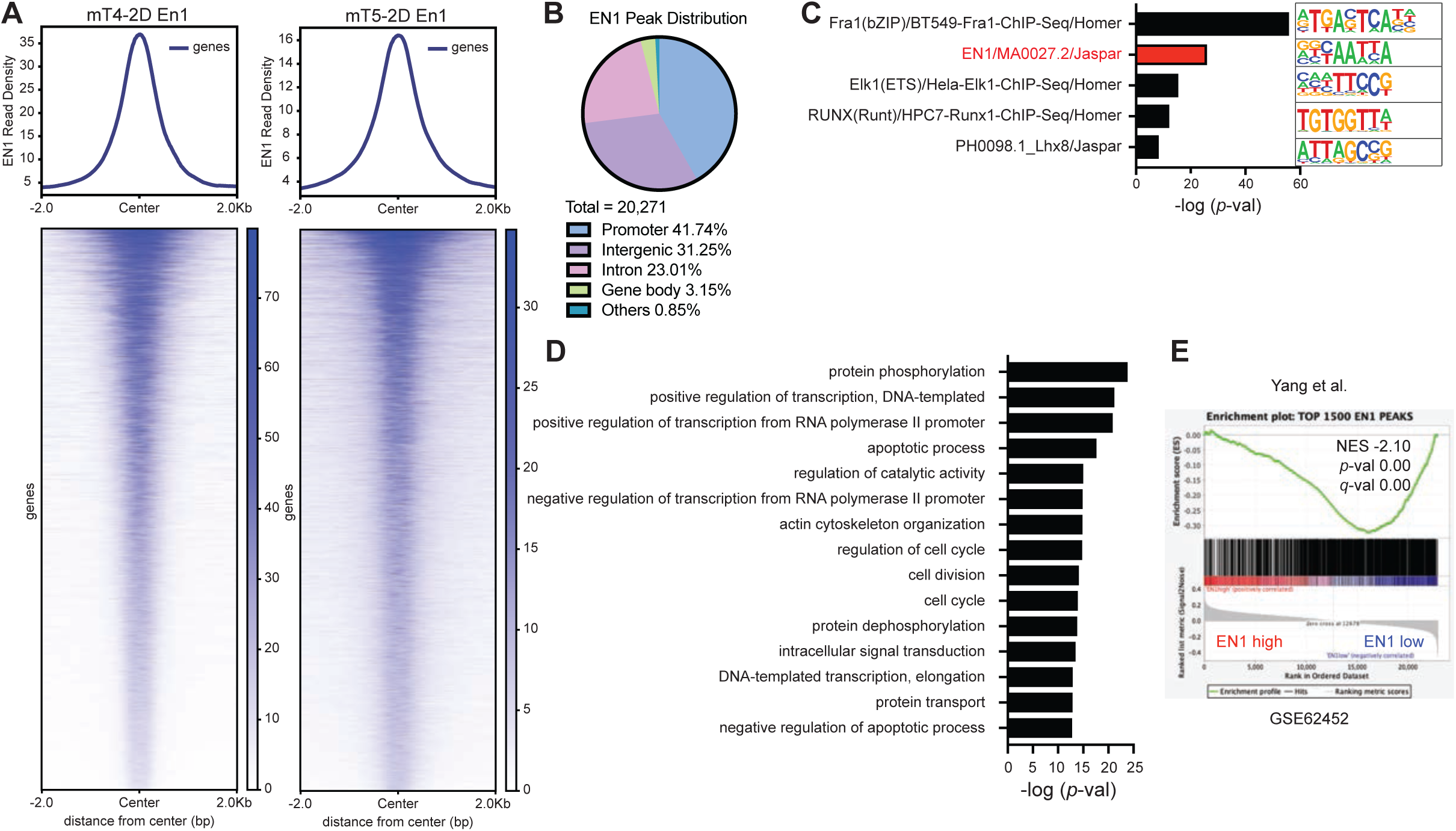
Identifying genomic targets of EN1 in murine PDA cells. (A) Density plots of CUT&RUN-seq signal of the EN1 DNA-binding peaks in mT4-2D and mT5- 2D cells with *FLAG-En1* cDNA. (B) Genome-wide distribution of the common EN1 peaks between mT4-2D *FLAG-EN1* and mT5-2D *FLAG-EN1* cells (n=20,271). (C) Homer motif analysis for the *de novo* motifs using the overlapping mT4-2D and mT5-2D EN1 peaks. (D) Gene ontology (GO) analysis using the Database for Annotation, Visualization, and Integrated Discovery (DAVID). The top 15 enriched pathways in biological functions were shown. (E) GSEA of the genes associated with En1 peaks in *EN1*-high vs. -low pancreatic cancer patients from Yang et al. (GSE62452). The genes associated with top 1,500 EN1 peaks among 20,271 common peaks were used for GSEA. NES, *p*-value, and FDR *q*-value were determined by GSEA.

### Identifying transcriptional targets of EN1 in PDA cells

Once EN1 genomic targets were identified, we next sought to pinpoint the transcriptional targets of EN1 in order to stratify if and/or how EN1 governs the expressions of its gene targets. To address this, we performed RNA-seq analysis of mM3P and mM15 organoids introduced with scramble or *En1*-targeting shRNAs (Fig. S3A). Differentially expressed gene analysis resulted in 154 differentially expressed genes (DEGs) with a statistical significance (*p*-val <0.05) (Fig. 5A). Of the total DEGs, 120 genes (79%) were upregulated upon *En1* knockdown, suggesting a transcriptional repressive role of EN1 in PDA. To then understand the functional significance of differentially expressed genes, we performed GO analysis using DAVID and GSEA (Fig. 5B-C). Both analyses showed the enrichment of apoptotic signaling pathways, in agreement with the functional annotations of EN1 genomic targets (Fig. 4D). Furthermore, hallmarks for E2F and MYC targets were also enriched in shScr organoids (Fig. 5C), well correlated with molecular signatures enriched in *EN1*-high patients from TCGA-PDDA dataset (Fig. 1J). To further elucidate the correlation between EN1 genomic and transcriptional targets, we performed GSEA and showed EN1 peak-associated genes were significantly enriched after *En1* knockdown (Fig. 5D), highlighting EN1 governs the gene expression predominantly through transcription repression. We also performed RNA-seq analysis for SUIT2 cells after *EN1* knockdown and identified 1,057 DEGs (Fig. S5A). Similar to murine cells, gene ontology analysis showed apoptosis, cell adhesion, and migration process were enriched in the DEGs (Fig. S5B), indicating the functional similarities of EN1 between murine and human PDA cells. Taken together, our data showed that as a TF, the major role of EN1 is transcription repression; in turn, the differentially expressed EN1 gene targets regulate anti-apoptotic, cell-cycle regulations, and cellular proliferation programs.

**Figure 5.**
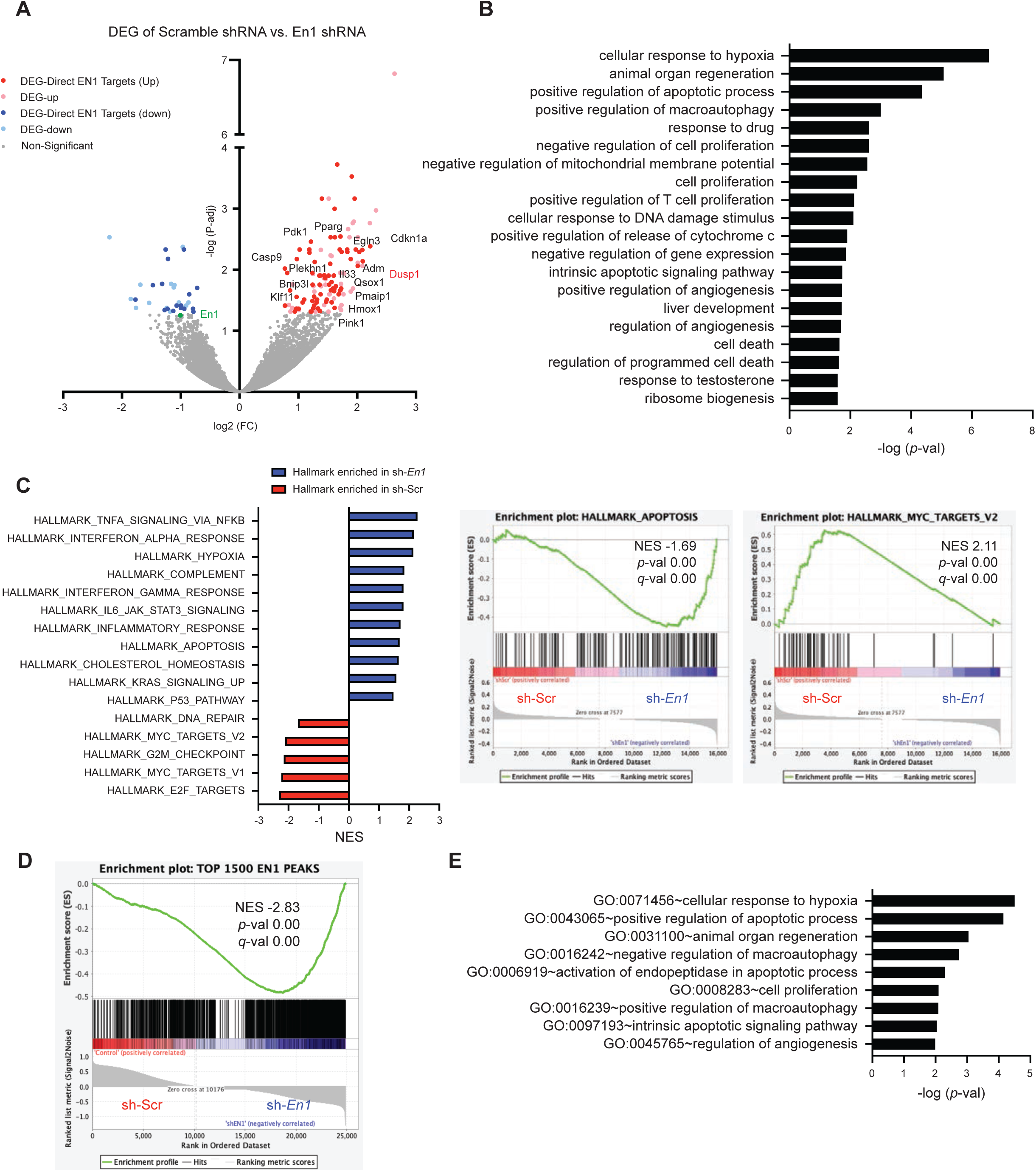
Identifying transcriptional targets of EN1 in PDA cells. (A) Volcano plot representing RNA-seq of mM3P and mM15 organoids with scramble (shScr) and two *En1* (sh*En1*) shRNA constructs. Differentially expressed gene analysis identified 154 differentially expressed genes (DEG). Among the DEGs, 120 genes were upregulated (red), and 32 genes were downregulated (blue) upon *En1* depletion; among which, 92 DEGs (dark red or dark blue) were the direct EN1 targets. DEG-direct EN1 target genes involved in cell death pathways were annotated. (B) GO analysis of the DEGs using DAVID. Top 20 significantly enriched biological functions were shown. (C) Normalized enrichment score (NES) of the GSEA Hallmark gene set in mM3P and mM15 organoids upon En1 knock-down. The top 16 significantly enriched hallmarks (left) and examples of the GSEA plots (right) are shown. Hallmark_Apoptosis and Hallmark_Myc_Targets_V2 gene sets were enriched in sh*En1* and shScr, respectively. (D) GSEA of the genes associated with top 1,500 EN1 peaks revealed that putative EN1 target genes were up-regulated upon *En1* depletion in mM organoids. (E) GO analysis of the EN1 direct target genes using DAVID. Among the genes associated with En1 peaks, commonly up-regulated genes upon En1 depletion were identified as direct targets of EN1.

### EN1 modulates gene promoter and enhancer activities to promote PDA progression

Our analysis of the EN1 binding regions in pancreatic cancer genome strongly suggested that EN1 targets promoters and enhancers through its DNA-binding domain. Since the majority of EN1 transcriptional targets were upregulated upon EN1 knockdown (Fig. 5A), we reasoned EN1 could repress gene transcription through altering promoter and enhancer activities. To better understand how EN1 regulates its target gene expression, we performed CUT&RUN-seq targeting active promoter marker tri-methylation of lysine 4 on histone H3 protein subunit (H3K4me3) and active enhancer marker acetylation of lysine 27 on histone H3 protein subunit (H3K27ac) using mT3-2D cell line overexpressing EN1. We then asked a question whether EN1 expression would alter H3K4me3 and H3K27ac occupancy in EN1 binding regions. H3K4me3 and H3K27ac CUT&RUN-seq analysis in mT3-2D cells revealed that H3K4me3 and H3K27ac occupancy were reduced surrounding EN1 binding sites (Fig. 6A) and the promoters of EN1 gene targets (Fig. 6B), indicating that EN1 binding reduced the activities of the target gene promoter and enhancer. Next, we performed H3K4me3 and H3K27ac CUT&RUN-seq with additional three biological replicates (mT4-2D, mT5-2D, and mT8-2D cell lines) upon EN1 overexpression and generated averaged meta-profiles (Fig. 6C-D). Similar with mT3-2D cells, upon EN1 overexpression, H3K4me3 and H3K27ac occupancies were decreased around EN1 binding regions and promoters of EN1 gene targets, suggesting EN1 can repress gene expression through modulating promoter and enhancer activities of its target genes. For instance, EN1 binds the promoter and distal enhancer of dual specificity phosphatase 1, *Dusp1* gene, and H3K4me3 and H3K27ac occupancies at these loci were reduced upon En1 overexpression (Fig. 6E), suggesting that EN1 could repress *Dusp1* expression through limiting the promoter and/or enhancer activities of *Dusp1.* DUSP1 is known to play a role in regulating cell death by dephosphorylating MAPKs (34, 35) and its expression was upregulated upon *En1* knockdown (Fig. 5A). To examine if EN1 affects ERK signaling activities, we performed phospho-ERK1/2 Western blotting in mM3P and mM15 organoids (Fig. 6F and S6A). En1 depletion resulted in decreased phospho-ERK1/2 signals, which was more pronounced in the reduced media (Fig. 6F and S6B), suggesting that EN1 positively regulates MAPK via repressing a negative regulator of MAPK pathway. Although EN1 genomic and transcriptomic targets are involved in cellular response to hypoxia and MYC pathways (Fig. 5), we did not observe any significant change in HIF-1α and c-MYC protein expressions upon *En1* knockdown (Fig. S6B). It has been shown that *En1* mutant mice shared a similar phenotype with *Ezh2* mutant mice (36, 37). Given the role of EZH2 in H3K27me3 (38) and the role of EN1 in transcriptional repression, we performed H3K27me3 CUT&RUN-seq with mT4-2D, mT5-2D, and mT8-2D cell lines upon EN1 overexpression. While we observed the enriched H3K27me3 occupancy at the known EZH2 binding regions (39), we saw negligible H3K27me3 occupancy at EN1 binding regions and no discernible changes upon EN1 overexpression (Fig. S6C), suggesting EN1-mediated transcription repression is independent of EZH2.

**Figure 6.**
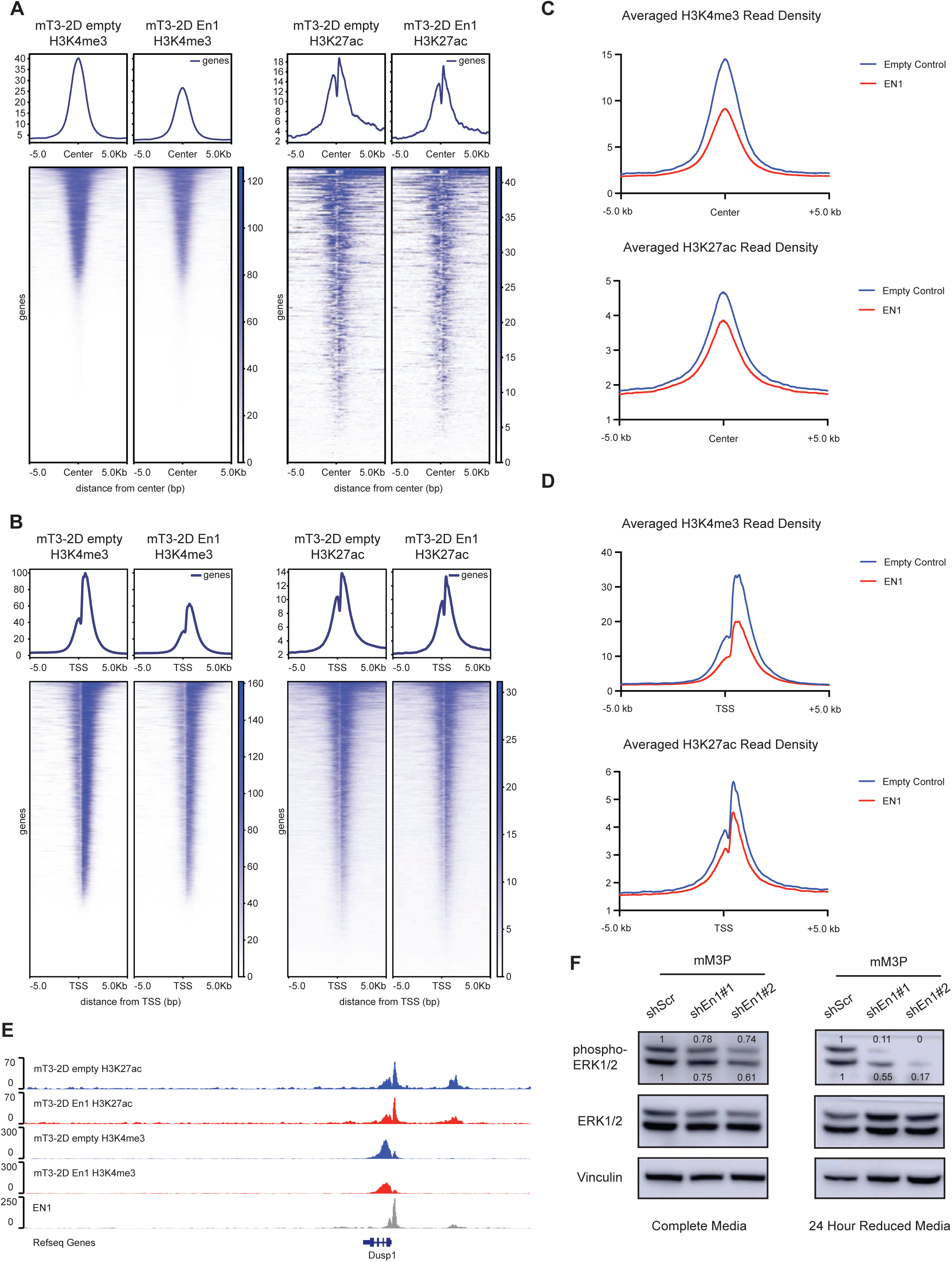
EN1 modulates gene promoter and enhancer activities to promote PDA progression. (A) Density plots of H3K4me3 (left) and H3K27ac (right) CUT&RUN-seq signals at EN1 genomic binding sites in mT3-2D empty | *FLAG-En1* cells. (B) Density plots of H3K4me3 (left) and H3K27ac (right) CUT&RUN-seq signals at EN1 peak- associated gene promoters and the transcription start sites (TSS) in mT3-2D empty | *FLAG-En1* cells. (C) Averaged density plots of H3K4me3 (left) and H3K27ac (right) CUT&RUN-seq signals at EN1 genomic binding sites in mT4-2D, mT5-2D, and mT8-2D empty | *FLAG-EN1* cells. (D) Averaged density plots of H3K4me3 (left) and H3K27ac (right) CUT&RUN-seq signals at EN1 peak-associated gene promoters and the TSS in mT4-2D, mT5-2D, and mT8-2D empty | *FLAG-EN1* cells. (E) Representative gene browser track of H3K27ac, H3K4me3, and EN1 CUT&RUN-seq signal at *Dusp1* gene in mT3-2D empty (blue) and *FLAG-EN1* (red) cells. (F) Western blot analysis to determine the protein expression of phospho-ERK1/2 (Thr202/Tyr204) and total ERK1/2 in mM3P organoids with scramble (shScr) and two independent *En1* (sh*En1*) shRNA constructs. Blots on the left showed organoids cultured in the complete organoid media and on the right showed organoids cultured in the reduced media for 24 hours before harvesting. Band intensity was determined by ImageJ.

### EN1 promotes PDA progression in GEMMs and PDA patients

Next, we asked whether EN1 deficiency in pancreatic epithelial cells could delay PDA progression in genetically modified mouse models (GEMMs). To this end, we crossed the conditional knock-out alleles of *En1* (aka *En1^flox/flox^*) with KPC mice to generate KPEC (*Kras^+/LSL-G12D^; Trp53^+/LSL-R172H^; En1^flox/flox^; Pdx1-Cre*) mice (Fig. 7A). There was no gross defect in pancreatic development when inactivating EN1 in the pancreas of EC mice (Fig. S7A). Long-term survival analysis showed that EN1 inactivation extended the animal overall survival (Fig. 7B), with the medium survival of 191 days for the KPEC mice and 125 days for the KPC mice. To illustrate the effect of *En1* inactivation in PDA progression, we sacrificed 10 mice per genotype at 120 days age for histopathological analysis. Histopathological analysis of KPC and KPEC mice showed the KPEC mice had significantly less percentage of abnormal pancreata, including acinar to ductal metaplasia (ADM), PanINs, and PDA, compared to KPC pancreata at 120 days of age (Fig. 7C). Of the examined animals at 120 days of age, 80% of KPC mice developed PDA compared to only 40% the KPEC mice that had developed PDA (Fig. 7D). One tumor-derived organoids (1 out of 4) from KPEC mice that we tested harbored unrecombined alleles of *En1* (Fig. S7B), suggesting that there might be a selective advantage for the unrecombined allele of *En1* during PDA progression of a certain KPEC mice. Overall, *En1* deficiency significantly attenuated PDA progression in our autochthonous mouse model. To confirm our findings in human PDA patient setting, we performed EN1 immunohistochemistry in the paired primary tumors and liver metastases tissue microarray from 19 PDA patients of the Rapid Autopsy Program (Fig. 7E). We found 7 out of 19 patients had a higher EN1 protein expression in the metastatic lesions compared to their paired primary tumors. Consistent with our finding that EN1 is a prognostic factor in PDA, EN1 protein expression level in the primary tumor was inversely correlated with the patient survival data (Fig. 7F). Taken together, our data showed that aberrant expression of EN1 facilitates PDA progression, resulting in poor survival of PDA GEMMs and patients.

**Figure 7.**
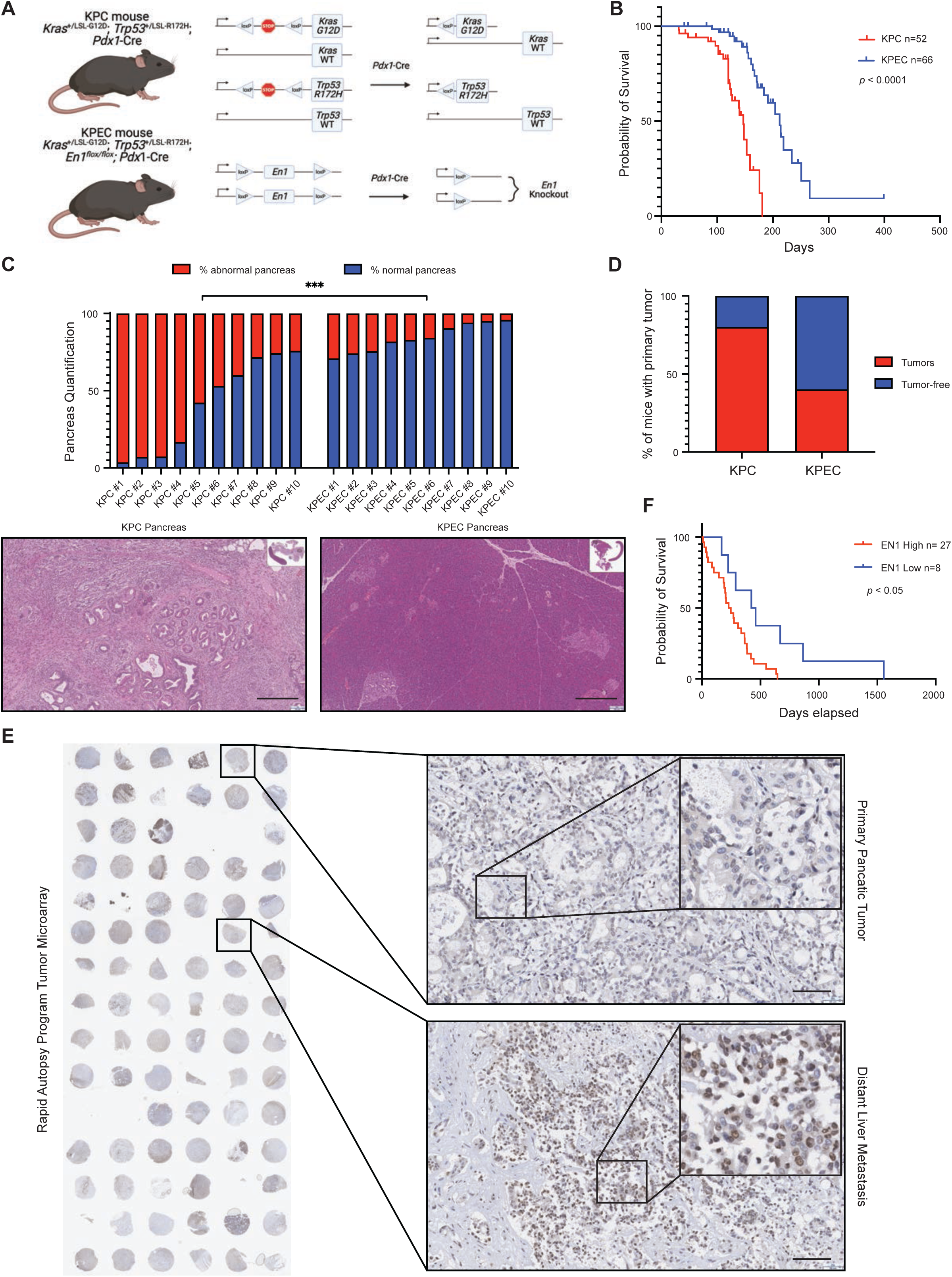
EN1 promotes PDA progression in genetically engineered mouse models and PDA patients. (A) Schematic representation of the genetically engineered mouse models with *Kras^+/LSL−G12D^, Trp53^+/LSL−R172H^, Pdx1-Cre* (KPC) and *En1^flox/flox^* (KPEC) alleles. (B) Kaplan-Meier plot of KPC (n = 52) and KPEC (n = 66) mice survival. The median survival of KPC mice is 147 days and the median survival of KPEC mice is 212 days. \*\*\*\**p*<0.0001 was determined by Log-rank (Mantel-Cox) test and GehIBreslow-Wilcoxon test. (C) Bar plot representing the percentage of abnormal pancreata (red) and normal pancreata (blue) from the KPC mice (n=10) and KPEC mice (n=10) at 120-day age. Representative H&E staining of KPC pancreas (bottom left, scale bar, 300 μm) and KPEC pancreas (bottom right, scale bar, 300 μm). (D) Quantification of the number of mice bearing tumors at 120-day age from KPC (n=10) and KPEC (n=10) mice. (E) Immunohistochemistry staining of EN1 in 19 human pancreatic and metastatic specimens from rapid autopsies (left). Representative image of a primary tumor EN1 IHC staining from patient #55 (Top right, scale bar, 100 μm). Representative image of a liver metastasis EN1 IHC staining from patient #55 (Bottom right, scale bar, 100 μm). (F) Kaplan-Meier plot of patient days survived after diagnosis corresponding to EN1-high (n=27) vs. -low (n=8) from the tissue microarray IHC. \**p*<0.05 was determined by Log-rank (Mantel-Cox) test and Gehan-Breslow-Wilcoxon test.

## Discussion

Non-mutational epigenetic reprogramming is one of the hallmarks of cancer (40). A growing body of evidence highlights critical roles of epigenetic alterations in carcinogenesis, including PDA. Previously, we and others have shown that aberrantly expressed transcription factors (e.g., TP63, FOXA1, EVI1, and TEAD2) alter pancreatic epigenome, thereby promoting PDA progression and a molecular subtype transition (4, 5, 7, 41, 42). Patients with metastatic PDA have a strikingly poor prognosis and limited response rate to current first line chemotherapies, including FOLFIRINOX and gemcitabine/nab-Paclitaxel (43, 44). The poor clinical outcome could be attributed by intrinsic chemoresistance of the cancer cell, or pro-survival program acquired during pancreatic carcinogenesis, which might be mediated through aberrant expressions of TFs and subsequent alteration in epigenetic landscapes and gene expressions. A better understanding of these mechanisms would allow us to identify potential targets and improve patient survival. Here, we identified aberrant expressions of EN1, a neuro-development TF in the late stage of PDA, resulting in enhancer reprogramming and endows aggressive characteristics in PDA progression. EN1 has been shown to be a pro-survival factor in brain development and associated with poor prognosis in multiple cancer types, such as adenoid cystic sarcoma, triple negative breast cancer, nasopharyngeal carcinoma, and osteosarcoma (12–18). Our data showed that EN1 perturbations altered the expression of a number of genes involved in apoptosis-, MYC-, hypoxia- and E2F-related pathways. For instance, we found that EN1 depletion altered MAPK pathways likely through the up-regulation of the negative regulators such as DUSP1, promoting cell survival. Collectively, EN1-mediated transcriptional alterations render the aggressive characteristics seen in our *in vitro* and *in vivo* studies. This observation highlights the critical role of developmental TF-mediated epigenetic reprogramming in cancer and might offer a unique therapeutic opportunity to exploit EN1-mediated epigenetic vulnerability in PDA.

While TFs are generally thought to be undruggable, it is possible to target the critical interacting proteins or functionally important downstream genes of the TFs. EN1 is known to function as a transcription repressor via the EH1 domain (45). Consistent with the known role as a transcriptional repressor, we showed a majority of EN1 target genes (79%) were upregulated upon EN1 depletion, suggesting that EN1 is predominantly a transcription repressor in the PDA context. The detailed molecular mechanisms of how EN1 reduced H3K27ac and H3K4me3 occupancy remain unknown and should be further explored. Thus, it would be worthwhile to identify repressive protein complexes that EN1 recruits to its genomic binding sites in PDA. For instance, in triple negative breast cancer (TNBC), the synthetic peptides targeting EN1 protein-protein interaction domains have been shown to induced cellular apoptotic responses *in vitro* (13). Likewise, targeting strategies of other interacting proteins in TNBC, such as TLE3, TRIM24-TRIM28-TRIM33 complex (18) and BRD4-S (46), might attenuate EN1-mediated aggressive cancer phenotypes in the breast cancer context. In addition, inhibition of direct EN1 downstream target genes might also be a novel therapeutic strategy. For example, *Dusp1*, a phosphatase negatively regulating ERK, JNK, and p38 MAPK activities (34), was identified as a direct repressive target of EN1 in our study. Thus, En1 depletion resulted in anti-survival phenotype, likely through up-regulation of DUSP1, a negative regulator of MAPK. A previous study has also shown that DUSP1 can antagonize a pro-survival signal upon gemcitabine treatment in PDA (47). Similarly, the downregulation of DUSP1 has been shown to confer pro-tumorigenic and metastatic characteristics (e.g., proliferation, migration, invasion, anti-apoptosis) in other cancer types, such as bladder and prostate cancers (48, 49). It should be noted that EN1 genomic binding sites appear to be context dependent since we did not find the EN1 target genes associated with WNT and Hedgehog signaling pathways that were previously identified in TNBC (16, 18). It is possible that the pre-existing epigenetic landscape in different cell types dictates the EN1 binding sites.

EN1 is an essential gene during embryonic development and its expression in the neuroepithelium is required to form midbrain and hindbrain (11). Within the adult central nervous systems, the mesodiencephalic dopaminergic neurons constitutively utilize EN1 to maintain the cellular identity, survival, outgrowth, and pathfinding (11). A line of evidence appears to point out that PDA exhibits neurodevelopment-related programs, such as axon guidance pathways, for their survival and tumorigenicity (9), while cancer cells generally utilize transcriptional programs associated with the cell-lineage for survival (8). In addition, a recent single-nucleus analysis of PDA samples identified a distinct neural-like progenitor (NRP) tumor cell type from patients that received the neoadjuvant therapies. Genes enriched in the NPR subtype were linked to axon guidance pathways, cell-cell adhesions, migrations, and negative regulations of cell death (50). Although EN1 was not differentially expressed in the NRP PDA subpopulation, the *EN1*- mediated transcriptional program, including axon guidance, cell-cell junction organizations, negative regulation of apoptosis, cell migration, and cytoskeleton organizations, may exert similar functions to the NRP-related programs in PDA as the neural-related genes were expressed within invasive epithelia of PDA to support cell survival and the development of therapeutic resistance. This observation highlights the clinical significance of aberrantly expressed TFs and their contributions to pancreatic epigenome, in turn promoting PDA progression, metastasis, and chemotherapeutic resistance.

In summary, we provided new evidence that EN1, a neurodevelopmental TF, could be aberrantly expressed in the late stage of PDA progression. EN1 can regulate a set of genes that govern pro-survival signals, contributing to metastatic characteristics of PDA. Importantly, we identified the direct targets of EN1 in PDA and elucidated the effect of EN1 in pancreatic cancer epigenome, which provides path to develop novel and exploitable drug targets in the future.

## Experimental Models

### Mouse models

All experiments were performed in accordance with the Institutional Animal Care and Use Committee (IACUC) of the University of California Davis and the NIH policies of the laboratory animal use. The behaviors and characterization of KPC (*Kras^+/LSL-G12D^; Trp53^+/LSL-R172H^; Pdx1-Cre*) alleles with the C57BL/6J strain have been described previously (7). 129S6/SvEvTac mouse harboring *En1^tm8.1Alj^* allele were purchased from the Jackson Laboratory (JAX stock #007918) and the allele were introduced into KPC mice through a series backcrosses to generate KPEC (*Kras^+/LSL-G12D^; Trp53^+/LSL-R172H^; En1^loxP/loxP^; Pdx1-Cre)* mice. For histological analysis, KPC and KPEC mice were sacrificed at 120 days of age. All animals were housed in the specific pathogen-free conditions and were regularly monitored by the veterinarians.

### Human specimens

Human tissue microarrays from 39 patients were obtained from the Rapid Autopsy Program at the University of Nebraska Medical Center. Written informed consent was obtained prior to tissue acquisition from all patients. Samples were assessed to be tumor and metastasis based on pathologist analysis.

### Tissue culture conditions

Murine pancreatic primary tumor organoids (mT3, mT6, mT19, and mT23), metastatic organoids (mM1, mM3, mM6, and mM10), and tumor 2D cell lines (mT3-2D, mT4-2D, mT5-2D, and mT8- 2D) from the tumor-bearing KPC mice were established and characterized previously (6, 7). Murine pancreatic organoid culture media contains Advanced DMEM/F-12 (Thermo Fisher 12634028), 10mM HEPES (Thermo Fisher 15630080), 1% Penicillin-Streptomycin (Thermo Fisher 15140122), 1% GlutaMAX Supplement (Thermo Fisher 35050061), 0.5μM A 83-01 (Fisher Scientific 29-391-0), 0.05μg/mL mEGF (Fisher Scientific PMG8043), 0.1μg/mL hFGF-10 (Pepro Tech 100-26), 0.01μM hGastrin I (Fisher Scientific 30-061), 0.1μg/mL mNoggin (Pepro Tech 250-38), 1.25mM *N*-Acetyl-L-cysteine (Millipore Sigma A9165), 10mM Nicotinamide (Millipore Sigma N0636), 1X B-27 Supplement (Fisher Scientific 17-504-044), and 1x RSPO1- conditioned medium. Murine 2D culture media contains DMEM (Corning 10-013-CV), 10% FBS (Gen Clone 25-550H), and 1% Penicillin-Streptomycin. Human PDA cell lines SUIT2 (Glow Biologics GBTC-1088B), CFPAC1 (ATCC CRL-1918), BxPC3 (ATCC CRL-1687), and PATU 8988s (Glow Biologics GBTC-0209H) were cultured with RPMI 1640 (Corning 10-040-CV), 10% FBS, and 1% Penicillin-Streptomycin.

### Next-generation sequencing

#### Cleavage under targets & release using nuclease (CUT&RUN) assay

CUT&RUN assay was performed according to the manufacturer’s instructions (Cell Signaling Technology CST 86652). Briefly, cells were trypsinized (Fisher Scientific 25-300-062) into single cells and counted using 0.4% Trypan Blue Stain (Thermo Fisher T10282) and Countess 3 FL Automated Cell Counter (Thermo Fisher). 250,000 cells were used for each reaction and input sample. CST CUT&RUN Protocol Section I.A. “Live Cell Preparation” was followed to precipitate histone marks and the Section I.B. “Fixed Cell Preparation” was followed to precipitate FLAG-tagged EN1. 1μL of anti-acetyl-Histone H3 (Lys27) antibody (CST 8173), 1μL of anti-tri-Methyl-Histone H3 (Lys4) antibody (CST 9751), or 1μL of anti-FLAG® M2 antibody (Thomas Scientific C986X12) was added to each reaction. Antibody incubation was carried at 4°C for 16 hours. 50 pg sample normalization spike-in DNA was added into each reaction during DNA digestion and diffusion. Fragmented DNA was purified using ChIP DNA Clean & Concentrator (ZYMO Research D5205). Bioruptor (Diagenode) was used to sonicate the input samples for 13 cycles (30 sec on/30 sec off at high amplitude). Bioinformatics pipeline was described previously (7).

#### RNA preparation for sequencing

For 2D cells, 70% confluent cells were trypsinized into single cells to yield 2x10^6^ cells. For organoids, 70% confluent organoids were trypsinized into single cells to yield 5x10^5^ cells. Cells were lysed with TRIzol reagent (Fisher Scientific 15-596-026) and RNA was collected per manufacturer’s instructions. Isolated RNA was treated with PureLink on-column DNase set (Thermo Fisher 12185010) and purified using PureLink RNA mini kit (Thermo Fisher 12183018A). Bioinformatics pipeline was described previously (7).

#### Library preparation and sequencing

Library preparation and sequencing for CUT&RUN and RNA were performed by Novogene Co., LTD (Beijing, China). Briefly, for CUT&RUN, sample quality control was performed prior to library construction. Then, the DNA fragments were end repaired, A-tailed, and ligated with illumina adapters. Following, the DNA library was filtered by size selection and PCR amplification. Quantified DNA libraries were pooled and sequenced using NovaSeq6000 PE150. Quality controls, including sequencing quality distribution, sequencing error rate distribution, ATCG base distribution, and adapter filtering were performed before raw data delivery. For RNA, sample quality control was performed prior to library construction. Then, mRNA was purified from the total RNA using polyT-oligo beads. After fragmentation, first strand cDNA was synthesized using random hexamers, and the second strand cDNA was synthesized using dTTP. Following, the cDNA was end repaired, A-tailed, ligated with illumina adapters, size selection, amplification, and purification. Quantified libraries were pooled and sequenced using NovaSeq6000 and paired-end reads were generated. Quality controls, including removing adapter, poly-N, and low quality reads, and Q20, Q30, and GC content calculations, were performed before raw data delivery.

### Protein and DNA-related experiments

#### Cloning

FLAG-tagged *En1* cDNA (Neo-FLAG-En1) was subcloned into MSCV-PGK-Neo-IRES-GFP (Neo-Empty) plasmid (Addgene 105505). FLAG-tagged *EN1* expression plasmid (MSCV-FLAG- EN1) was obtained from VectorBuilder (Vector ID: VB220501-1183hep) and the negative control plasmid (MSCV-Empty) was generated using restriction enzyme AvrII (NEB R0174S) and EcoRI-HF (NEB R3101S), and blunting & ligation kit (NEB E0542S) to remove FLAG-EN1 sequence. *En1* and *EN1* shRNAs were obtained from the TRC shRNA library available at the Broad Institute (sh*En1* #1 TRCN0000082149, sh*En1* #2 TRCN0000414478, sh*EN1* #1 TRCN0000013899, sh*EN1* #2 TRCN0000013968) in pLKO.1 puro construct.

#### Reverse transcription and quantitative PCR

Total RNA was extracted using TRIzol reagents per manufacturer’s instructions as described in the RNA preparation for sequencing. RNA concentration was measured using NanoDrop 1000 (Thermo Fisher). 1500 ng of RNA was used for cDNA synthesis with high-capacity cDNA reverse transcription kit (Thermo Fisher 4368814). 1 μL of the cDNA was used for qPCR with *Power* SYBR green PCR master mix (Thermo Fisher 4368702) on LightCycler 480 instrument II (Roche Diagnostics). The qPCR results were quantified using the 2^(delta)(delta)Ct method with housekeeping gene *GAPDH*, *Gapdh*, or *ACTB*, *Actb* for data normalization. qPCR primer sequences used in the manuscript are listed in the Table S1.

#### Western blot analysis

70% confluent cells were harvested and lysed with protein extraction buffer (50 mM Tris pH 7.4, 1 mM EDTA, 150 mM NaCl, 1% NP-40, and 1x Halt protease inhibitor cocktail (Thermo Fisher 78437)) on ice for 30 minutes, centrifuged at 20,000 RCF 4°C for 20 minutes, and collected the supernatant. Protein was measured for concentration with protein assay kit (Bio-Rad 5000111) and denatured with sample reducing agent (Thermo Fisher NP0009). 10 μg protein lysate was loaded into 4 to 12% Bis-Tris 1.0 cm gels (Thermo Fisher NP0321BOX or NP0322BOX) and electrophoresis was carried using mini gel tank (Thermo Fisher A25977) at 120 V. Protein transfer to PVDF membrane (Millipore Sigma IPVH00010) was carried using transfer cell (Bio-rad 1703930) at 400 mA for 2 hours at 4°C. The membrane was blocked with 5% Non-fat milk dissolved in PBS with 1% Tween-20 (PBST) at room temperature for 1 hour, washed with PBST four times 5 minutes each, and incubated with diluted primary antibody at 4°C for 16 hours. The membrane was washed with PBST four times 5 minutes each and incubated with diluted secondary antibody at room temperature for 1 hour followed by PBST wash four times 5 minutes each. Luminol signals were developed using pico chemiluminescent substrate (Thermo Fisher 34577) and detected using Amersham Imager 600 (GE Healthcare Life Sciences). Antibody used in the manuscript are listed in the Table S2. For data analysis, phospho-ERK data normalization = (phospho-ERK bands intensities)/(ERK bands intensities/Vinculin bands intensities). HIF-1α and c-MYC data normalization = HIF-1α or c-MYC bands intensities/Vinculin bands intensities. Secondary normalization was analyzed by comparing the *En1* knockdown groups with the scramble control.

#### Genotyping

Mice toes were clipped at day 10.5 and the genomic DNA was isolated using 30μL TaqAN buffer (10mM Tris-HCl, 50mM KCl, 2.5 mM MgCl_2_, 0.45% NP-40, 0.45% Tween-20, and 3μL/mL of Proteinase K (NEB P8107S)) at 56°C for 1 hour followed by denaturation at 96°C for 10 minutes. *Taq* DNA polymerase (NEB M0273E) was used to PCR *Trp53* and *Cre*. Platinum hot start PCR master mix (Thermo Fisher 13000012) was used to PCR *Kras* and *En1*. PCR conditions for *Trp53, Trp53* het/homo, and *Cre* was 94°C 3 min, 40 cycles of 94°C 1 min / 60°C 1 min / 72°C 1 min, and 72°C 3 min. PCR conditions for *Kras* was 94°C 3 min, 35 cycles of 94°C 1 min / 69°C 2 min / 72°C 1 min, and 72°C 3 min. PCR conditions for *En1* was 94°C 2 min, 40 cycles of 94°C 30 sec / 60°C 30 sec / 72°C 30 sec, and 72°C 2 min. AmpliTaq Gold 360 master mix (Thermo Fisher 4398876) was used to PCR 1 loxP *En1*, and the PCR condition was 95°C 5 min, 40 cycles of 95°C 30 sec / 61°C 30 sec / 72°C 30 sec, and 72°C 5 min. PCR primer sequences used in the manuscript are listed in the Table S1.

#### Retrovirus production and infection

Retrovirus was produced in either Phoenix-AMPHO (ATCC CRL-3213) or Phoenix-ECO (ATCC CRL-3214) and lentivirus was produced in HEK293T (ATCC CRL-3216) via X-tremeGENE9 (Millipore Sigma 6365809001) transfection. Cells were first grown to 70% confluence in 10-cm tissue culture plate (Genesee Scientific 25-202). Before transfection, culture media was replaced with DMEM supplemented with 10% FBS. For retrovirus, 10μg transfer plasmid and 15μL X-tremeGENE9 reagent was mixed well in 400μL DMEM. For lentivirus, 5μg transfer plasmid, 2.25μg psPAX2 (Addgene 12260), and 0.75μg pMD2.G (Addgene 12259) was mixed well in 400μL DMEM. The mixture was incubated at room temperature for 20 minutes then added to the cell culture dropwise. The condition media was collected after 48-72 hours and filtered through a 0.2μm filter (PALL 4612). Organoid infection procedures were described previously (7). For infecting 2D cells, filtered conditional media was added to host cells grown at 50% confluence with 10μg/mL polybrene (Thomas Scientific C788D57 (EA/1)). Three days after infection, the cells were selected by 2μg/mL puromycin (Fisher Scientific 53-79-2), 1mg/mL Geneticin G418, or fluorescence-activated cell sorting (Sony SH800S).

### Histology

#### Immunohistochemistry staining

Paraffin-embedded tissue sections were first placed in an oven at 60°C for 30 minutes. The slides were placed in Histo-Clear (National Diagnostics HS-200) for two changes 10 minutes each, 100% EtOH two changes 2 minutes each, 95% EtOH two changes 2 minutes each, 85% EtOH two changes 2 minutes each, 75% EtOH two changes 2 minutes each, deionized distilled water (ddH_2_O) one change for 1 minute, and PBS one change for 1 minute. To retrieve antigens, the slides were placed in Citrate-EDTA buffer (Abcam ab93678), boiled using an electric pressure cooker (Cuisinart) for 10 minutes at low pressure, and slowly cooled at room temperature for 1 hour. Incubate the sections with PBS two changes 5 minutes each. The sections were then incubated with BLOXALL (Vector Lab SP-6000-100) for 10 minutes and 2.5% horse serum (Vector Lab S-2012-50) for 30 minutes. The sections were incubated with the horse serum diluted antibody for 16 hours at 4°C. Then, the slides were washed with PBST one change for 3 minutes and PBS one change for 3 minutes. Hereafter, the sections were processed using VECTASTAIN Universal ABC-HRP kit (Vector Lab PK-7200), DAB substrate kit (Vector Lab SK-4100), hematoxylin counterstain (Vector Lab H-3401), and mount with VectaMount (Vector Lab H-5000-60) per manufacturer’s instructions. Antibody used in the manuscript are listed in the Table S2. For hematoxylin and esosin (H&E) staining, paraffin-embedded tissue sections were first placed in Histo-clear for three charges 3 minutes each. Hematoxylin and Eosin stain kit (Vector Lab H-3502) then was used to stain for H&E per manufacturer’s instructions.

### In vitro assays

#### Colony formation assay

All cell lines were trypsinized to generate single cell suspensions and counted three times to average the cell counts. KPC-2D cell lines (1,000 cells) were resuspended in DMEM supplemented with 10% FBS and 1% Penicillin-Streptomycin and plated in 6-well tissue culture plates (Celltreat 229105) for 5 days. CFPAC1 (500 cells) and PaTu8988S (1000 cells) were resuspended in RPMI 1640 supplemented with 10% FBS and 1% Penicillin-Streptomycin and plated in 6-well tissue culture plates for 7 and 14 days respectively. SUIT2 (500 cells) and BxPC3 (1000 cells) were resuspended in RPMI 1640 supplemented with 10% FBS and 1% Penicillin-Streptomycin and plated in 24-well tissue culture plates (Corning 3527) for 5 and 14 days respectively. Colonies were stained at room temperature for 1 hour with 2% crystal violet (Thomas Scientific 30430001-1) diluted in 100% methanol to reach the 0.5% final concentration followed by tap water wash three times and running water wash for five minutes. The plates were imaged with a printer scanner (HP LaserJet Pro) and clonogenic growth was analyzed by ImageJ.

#### Tumor spheroid formation assay

All cell lines were trypsinized to generate single cell suspensions and counted three times to average the cell counts. 500 cells of CFPAC1 or SUIT2, 1000 cells of PaTu8988S or BxPC3, and 25,000 cells of KPC-2D cells were resuspended in 3D Tumorsphere Medium XF (PromoCell C-28075) and plated in ultra-low attachment 24-well plates (Millipore Sigma CLS3473-24EA) for 7 days. Culture suspensions were mixed well prior for imaging using EVOS M5000 imaging system (Fisher Scientific) under 4x bright field. Spheroids were analyzed by ImageJ.

#### Wound-healing assay

Cells were grown to 90% in 6-well tissue culture plates and wounded linearly using a 200 μL tip followed by three washes of PBS. 24 hours after, cell migration was imaged under 4x bright field. Percentage of migration was determined by ImageJ.

#### Boyden chamber invasion assay

Matrigel (Corning 356231) was first diluted in DMEM at 1:3 dilution. 100 μL diluted Matrigel was then placed in transwell insert (Neta Scientific SIAL-CLS3464) and incubated in the tissue culture incubator for 3 hours. 600 μL of DMEM supplemented with 10% FBS was added to the lower chamber. 50,000 per 200 μL of cells were then added on top of the solidified Matrigel and incubated for 24 hours. After, the transwell was removed and gently scrubbed with a cotton swab and washed twice with PBS. Cells were then stained with SYTO 13 GFP nucleic acid stain (Life Technologies S7575) per manufacturer’s instructions and imaged under 4x GFP channel for cell count.

#### Organoid survival assay

Pancreatic organoids were maintained in the complete organoid media prior to single cell dissociation as previously described (7). 5,000 cells were resuspended in 50 μL Matrigel and plated into a 24-well tissue culture plate for 4 or 5 days. Organoids were cultured either in the organoid complete media and the reduced media (DMEM supplemented with 10% FBS and 1% Penicillin-Streptomycin). Organoids were imaged under 4x bright field and quantified by ImageJ.

#### Organotypic tumor-on-a-chip assay

Detailed protocols to micro-fabricate tumor-blood vessel is described previously (28). Briefly, the organotypic PDA on-a-chip was made with polydimethysiloxane gaskets and coated with 0.1 mg/mL poly-L-lysine (Millipore Sigma 4707), 1% glutaraldehyde (Electron Microscopy Sciences 16310), and 2.5 mg/mL rat tail collagen I (Corning 354236). Mouse KPC mT3-2D empty and En1 cells were grown in DMEM (Corning 10-013-CV) and human umbilical vein endothelial cells in EGM-2 (Lonza CC-3162). PDA cells were seeded in day 1, and endothelial cells were seeded in day 2. Media in the PDA channel and biomimetic blood vessel was refreshed and monitored daily through the experiment.

#### In vivo assays

Female 6- to 8-week-old syngeneic C57BL/6J or athymic immune-compromised (NU/NU) nude mice were purchased from the Jackson Laboratory (000664) and Charles River Laboratory (088), respectively. All animal procedures were conducted in accordance with the IACUC at Cold Spring Harbor Laboratory. For subcutaneous transplantation, mice were first anesthetized by isoflurane. 500,000 cells resuspended with 50μL Matrigel were injected into the left flank of subcutaneous space. For orthotopic transplantation, mice were first anesthetized by isoflurane. Iodine solution was applied to the incision site. Then, a small incision (∼1 cm) was made at the upper left quadrant of the abdomen. Following, 500,000 cells resuspended with 50μL Matrigel was injected into the pancreas parenchyma. For tail vein transplantation, restrained mice were injected with 50,000 cells resuspended in 50μL PBS intravenously through the tail vein.

#### Data Availability

The sequencing data that support the findings of this study are openly available in NCBI Gene Expression Omnibus database: GSE228805.

**Figure S1. Identification of EN1 in organoid survival assay and its association with the PDA aggressive phenotype.**

(A) Image-based quantification of mT and paired mM organoid survival in the reduced media 4 days post-cell seeding. n=3, mean ± SEM.

(B) Bar plot representing the number of passages the organoids underwent. Arrow indicating the organoids can be passaged continuously in the reduced media.

(C-D) Image representation of the development of organoid survival assay in the indicated mT and mM organoid pairs with *Trp53* knockout and *Rosa26* knockout control in the reduced media.

Scale bar, 1mm

(E-F) Subcutaneous transplantation of mT6 organoids with *Trp53* knockout using CRISPR/Cas9. gRNA against *Rosa26* locus was used as a control. The tumors were imaged (E) and quantified for tumor weight (F) at 39 days post-injection. n=5 per group, mean ± SD. Scale bar, 10mm.

(G-H) After two gRNAs targeting wild-type *Trp53* were introduced in mT organoids (parental organoids), mT organoids were subjected to the reduced media in order to enrich p53 LOH mT organoids. These organoids (n = 3 for *Rosa26* gRNA, n = 3 for parental organoids and n = 6 for the enriched population for *p53* LOH) were subcutaneously transplanted in NSG mice. The tumors were collected 5 weeks post-injection (G) and quantified by tumor weight (I). Scale bar, 10 mm.

(I) *Batf2*, *Foxa1*, *Gata5*, *Prrx2*, *Pax9*, and *Trerf1* mRNA expression in mM organoids relative to mT organoids from Oni et al. (GSE63348) and Roe et al. (GSE99311). Each dot represents an organoid line.

(J-K) Relative cell density in the organoid survival assay from Figure 1C (J) and Figure 1D (K) was quantified. Mean ± SEM is shown.

(L) EN1 IHC of the pancreatic primary tumor, lung and peritoneal metastases from a KPC mouse. Scale bar, 100 µm.

(M) *EN1* normalized read count of primary purified circulating tumor cells from pancreatic cancer patients (localized and metastatic) and healthy donors from Franses et al. (GSE144561).

(N) *EN1* mRNA mean counts per cell of 24 pancreatic patients from Peng et al. scRNA-seq (CRA001160). Mean ± SD is shown.

(O) Squamous subtype markers, *TP63* and *KRT5* mRNA counts in *EN1*-high (T6, T14, T16, T17) vs. -low (T1, T2, T3, T5) PDA patients from Peng et al. scRNA-seq (CRA001160).

Unless otherwise indicated, *p*-values were determined by unpaired student’s *t* test (two-tail) and *, **, ***, **** indicate *p*-val < 0.05, < 0.01, <0.001, <0.0001, respectively.

**Figure S2. Gain-of-function experiments revealed that EN1 promotes PDA metastatic properties.**

(A) Relative *En1* mRNA expression determined by RT-qPCR in mT3-2D and mT23-2D cell lines with (*En1*) and without (empty) *En1* cDNA overexpression.

(B) mT3-2D and mT23-2D cells with *En1* cDNA were subjected to cell proliferation assay compared to empty vector control. Cell proliferation rate was determined by ATP-based cell viability assay using CellTiterGlo and luminescence was measured daily for 4 days and normalized to day 1. n=3 per time point, mean ± SD.

(C) mT23-2D with *En1* cDNA were subjected to wound-healing assay compared to empty vector control, and the area of the closed wound was quantified at 0- and 24-hour post-scratching, and the percentage of wound closure was calculated. n=3, mean ± SD.

(D) Relative *EN1* mRNA expression in human PDA cell lines determined by RT-qPCR. n=3, mean ± SD.

(E) Western blot analysis to determine the protein expression of FLAG-tagged EN1 compared to the plasmid without *EN1* cDNA (empty) control in CFPAC1 and PaTu 8988s cell lines.

(F) PaTu8988s empty and *EN1* cells were subjected for colony formation assay for 14 days, and the colonies were stained by crystal violet (right) and quantified (left) by percentage growth area. n=3, mean ± SD.

(G) PaTu8988s empty and *EN1* cells were subjected for anchorage-independent tumor spheroid formation assay for 7 days, and the numbers of spheroids were monitored (right) and quantified (left). n=3, mean ± SEM. Scale bars, 350 μm.

Unless otherwise indicated, *p*-values were determined by unpaired student’s *t* test (two-tail), and * and *** indicate *p*-val < 0.05 and <0.001, respectively.

**Figure S3. EN1 is necessary to acquire the metastatic characteristics of PDA.**

(A) Relative *En1* mRNA expressions determined by RT-qPCR in mM3P, mM15, and mM10 organoids with shRNA targeting *En1* mRNA coding region (sh*En1* #1 CDS) and *En1* mRNA 3’ untranslated region (sh*En1* #2 3’UTR) compared to the scramble shRNA (sh-scr) control organoid. n=3, mean ± SD.

(B) Image-based quantification of organoid growth for mM3P and mM10 shScr, sh*En1* #1, and sh*En1* #2 organoids in the complete media. Organoid growth was normalized to day 1. n=3, mean ± SD.

(C) shScr and sh*En1* mM10 organoids were subjected to organoid survival assay for 4 days (top) and quantification of organoids (bottom). Scale bars, 1mm.

(D) shScr and sh*En1* mM10 organoids were subjected to colony formation assay for 7 days, and the colonies were stained by crystal violet (left) and quantified (right) by percentage growth area.

(E) Summary of metastases from mM3P shScr (n=5) and sh-*En1* orthotopic transplants (n=5), 7 weeks post-transplantation. Depletion of En1 reduced liver metastasis frequency. Fisher’s exact test, *p*-val <0.05.

(F) Representative H&E staining of liver metastasis from the orthotopic injections of mM3P shScr and sh*En1* organoids. Scale bar, 50 µm.

(G) Representative H&E staining of lung metastasis from the orthotopic injections of mM3P shScr and sh*En1* organoids. Scale bar, 50 µm.

(H) Relative EN1 mRNA expressions determine by RT-qPCR in SUIT2 (left) and BxPC3 (right) cell lines with shRNAs targeting *EN1* mRNA compared to the scramble shRNA. n=3, mean ± SD.

(I) shScr and sh*EN1* BxPC3 cells were subjected to colony formation assay for 2 weeks, and the colonies were stained with crystal violet (top) and quantified (bottom) for the percentage growth area. n=3, mean ± SD.

(J) shScr and sh*EN1* BxPC3 cells were subjected to anchorage-independent tumor spheroid formation assay for 7 days, and the numbers of spheroids were monitored (left) and quantified (right). n=3, mean ± SD. Scale bars, 350 μm.

Unless otherwise indicated, *p*-values were determined by unpaired student’s *t* test (two-tail) and *, **, ***, **** indicate *p*-val < 0.05, < 0.01, <0.001, <0.0001, respectively.

**Figure S4. Characterization of EN1 binding regions in PDA genome.**

(A) Western blot analysis to determine the protein expression of FLAG-tagged EN1 compared to the empty control in mT3-2D, mT4-2D, mT5-2D, mT8-2D, and mT19-2D cells.

(B) Homer motif analysis for the known motifs using the overlapping mT4-2D and mT5-2D EN1 peaks. *p*-value was determined by HOMER.

(D) Genomic Regions Enrichment of Annotations Tool (GREAT) analysis of the overlapping mT4-2D and mT5-2D EN1 peaks showing the top 15 enriched pathways in biological functions. *p*-value was determined by GREAT.

(C) Overlapping EN1 peaks identified in triple-negative breast cancer (TNBC) cells (MCF7, SUM149, and SUM159) and mT4-2D & mT5-2D cells. To properly compare murine and human genomes, we convert the EN1 peak positions identified in mT-2D cell lines (NCBI37/mm9) to human genome assembly (GRCh37/hg19) using UCSC Lift Genome Annotations tool. Of the 20,271 EN1 peaks identified in mT-2D cell lines, 17,967 peaks were LiftOver successfully. Among the TNBC cell lines, 182, 213, and 471 peak overlaps were identified in MCF7, SUM159, and SUM149 cells, respectively.

(E) GSEA of the top 1500 EN1 peak-associated genes in *EN1*-high vs. -low pancreatic cancer patients, organoids, or cell lines from Tiriac et al. (phs001611.v1.p1), Klett et al. (GSE101448), Jiang et al. (GSE71989), TCGA-PAAD, and Puleo et al. NES, *p*-value, and FDR *q*-value were determined by GSEA.

**Figure S5. EN1 genomic targets were up-regulated upon EN1 depletion.**

(A) Volcano plot representing RNA-sequencing of mM3P and mM15 organoids with scramble (shScr) and two *En1* (sh*En1*) shRNA constructs identified 154 differentially expressed genes (DEG). Among the DEGs, 120 genes were upregulated, and 32 genes were downregulated after *En1* knockdown. Blue dots represent EN1 genomic targets and red dots represent non-genomic targets of EN1.

(B) Volcano plot representing RNA-sequencing of SUIT2 shScr and two independent sh*EN1* constructs identified 1056 differentially expressed genes (DEG). Among the DEGs, 638 genes were upregulated, and 418 genes were downregulated after *En1* knockdown.

(C) DAVID analysis of the DEGs after *EN1* knockdown in SUIT2 cells showing the top 10 enriched pathways in biology functions. *p*-value was determined by DAVID.

**Figure S6. EN1 regulates MAPK pathways but not MYC or HIF-1α-dependent hypoxia responses and EN1-mediated transcription repression is independent of EZH2.**

(A) Western blot analysis to determine the protein expression of phospho-ERK1/2 (Thr202/Tyr204) and total ERK1/2 in mM15 organoids cultured in the complete media with scramble (shScr) and two independent *En1* (sh*En1*) shRNA constructs.

(B) Western blot analysis to determine the protein expression of phospho-ERK1/2 (Thr202/Tyr204) and total ERK1/2 in mM3P and mM15 organoids cultured in the reduced media for 24 hours with scramble (shScr) and two independent *En1* (sh*En1*) shRNA constructs. Vinculin data for mM3P was duplicated as Figure 6F right panel. Band intensity was determined by ImageJ.

(C) Averaged density plots of H3K27me3 CUT&RUN-seq signals around EZH2 genomic binding sites (left) and EN1 genomic binding sites (right) in mT4-2D, mT5-2D, and mT8-2D empty | *FLAG-EN1* cells.

**Figure S7. *En1* knockout does not affect the development of murine pancreas and Cre recombination is not prevalent in all animals.**

(A) H&E staining of pancreas isolated from *En1^flox/flox^; Pdx1-Cre* (EC) mouse at 67 days age.

(B) PCR analysis of 1 loxP-*En1* (recombined, top) and LSL*-En1* cassette (unrecombined, bottom) using tumor and metastasis organoids derived from KPEC mice. T: tumor; M: metastasis; L: liver; and P: peritoneum. Expected En1 1 loxP size: 600 bp; expected LSL-En1 size: 380 bp.

## Acknowledgements

We would like to thank all members of the D.A.T. and C.-I.H. laboratories for helpful discussions and suggestions throughout the course of this study. The authors would like to thank the Cold Spring Harbor Cancer Center Support grant (CCSG, P30CA045508-29) shared resources: DNA Sequencing, Animal facility and the Histology core, as well as the UC Davis, The Center for Genomic Pathology Laboratory and Qian Chen for histology, and DNA sequencing core and Shelly Williams for DNA sequencing. The authors would like to thank Sarah Wang and Suyakarn Archasappawat for mouse colony management; Qi Tian, Abigail Brand, Shounak Ranabhor, Keely Ji, Fatimah Al-Musawi, Neha Ramesh, Shou Kitahara, Tha Thu, Madison Hall, Cynthia Huang, Ayushi Borthakur, Clarissa Im, Nicholas Kaiser, Sriansh Pasumarthi, and Jamie Lee for mouse genotyping. We also thank Omar Younis for histology and Keely Ji for providing her feedback. J.X. is supported by the NIEHS-funded predoctoral fellowship (T32 ES007059). C.-I.H. is supported by the National Cancer Institute (NCI) K22CA226037 and R37CA249007, and the UC Davis Comprehensive Cancer Center Pilot Grant (NCI P30CA093373). D.A.T. is a distinguished scholar of the Lustgarten Foundation and Director of the Lustgarten Foundation– designated Laboratory of Pancreatic Cancer Research. D.A.T. is also supported by the Cold Spring Harbor Laboratory Association, the V Foundation, the Thompson Foundation, and the NIH (NIH P30CA45508, U01CA224013, U01CA210240, and R01CA188134). D.A.T. is supported by the Simons Foundation (552716). Y.P. is supported by the NCI R50CA211506.

