## Supplemental Figures for "*Engrailed-1* Promotes Pancreatic Cancer Metastasis"

### Figure S1

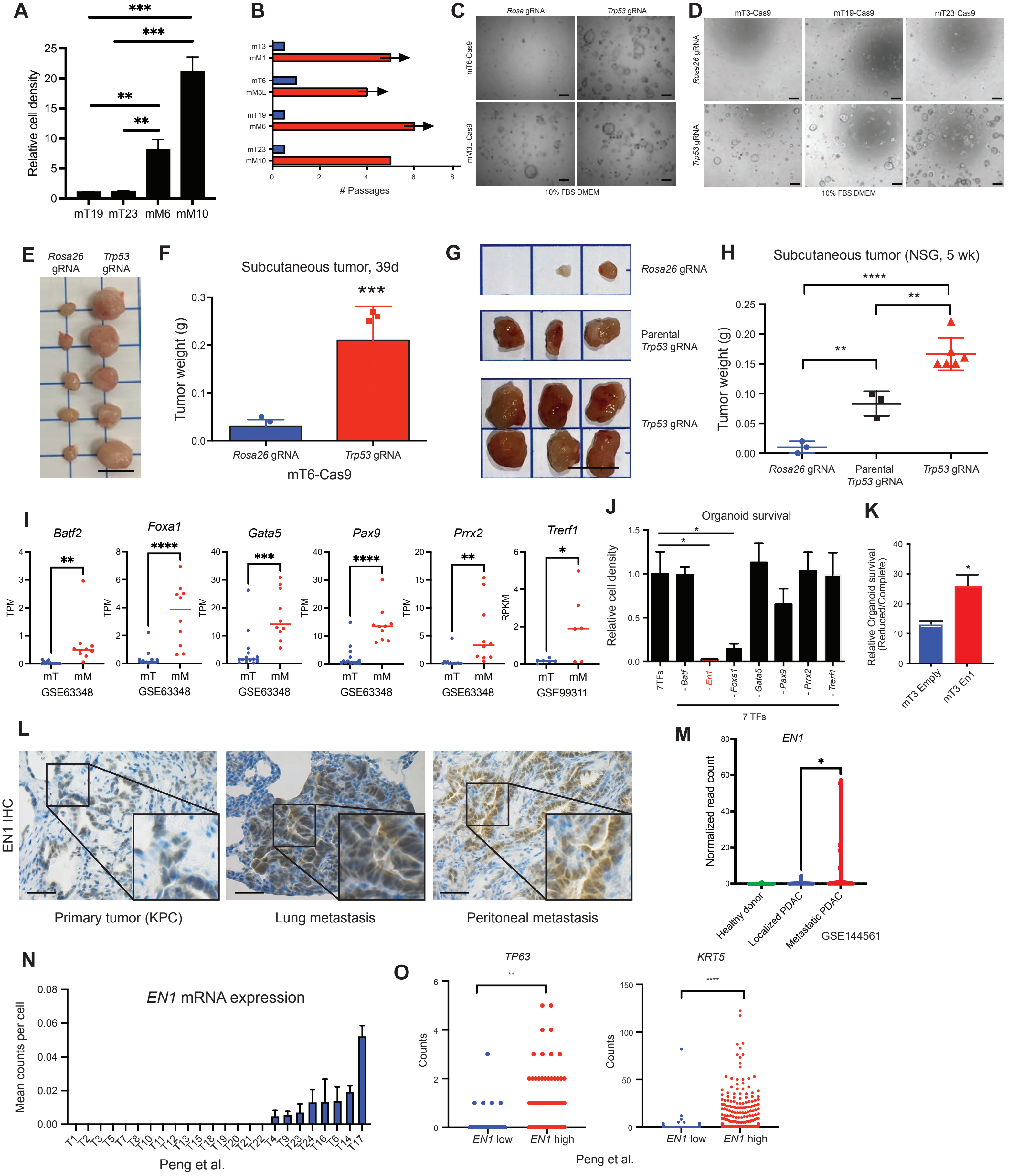
A
Relative cell density
mT19 mT23 mM6 mM10
B
mT3 mM1 mT6 mM3L mT19 mM6 mT23 mM10
### Passages
C
Rosa gRNA
Trp53 gRNA
mT6-Cas9
mM3L-Cas9
10% FBS DMEM
D
mT3-Cas9
mT19-Cas9
mT23-Cas9
Rosa26 gRNA
Trp53 gRNA
10% FBS DMEM
E
Rosa26 gRNA
Trp53 gRNA
F
Subcutaneous tumor, 39d
Tumor weight (g)
Rosa26 gRNA
Trp53 gRNA
mT6-Cas9
G
Rosa26 gRNA
Parental Trp53 gRNA
Trp53 gRNA
H
Subcutaneous tumor (NSG, 5 wk)
Tumor weight (g)
Rosa26 gRNA
Parental Trp53 gRNA
Trp53 gRNA
I
Batf2
Foxa1
Gata5
Pax9
Prrx2
Trerf1
TPM
RPKM
mT mM
GSE63348
GSE99311
J
Organoid survival
Relative cell density
7TFs
- Batf
- En1
- Foxa1
- Gata5
- Pax9
- Prrx2
- Trerf1
7 TFs
K
Relative Organoid survival (Reduced/Complete)
mT3 Empty
mT3 En1
L
EN1 IHC
Primary tumor (KPC)
Lung metastasis
Peritoneal metastasis
M
EN1
Normalized read count
Healthy donor
Localized PDAC
Metastatic PDAC
GSE144561
N
EN1 mRNA expression
Mean counts per cell
T1 T2 T3 T5 T7 T8 T10 T11 T12 T13 T15 T18 T19 T20 T21 T22 T4 T9 T23 T24 T16 T6 T14 T17
Peng et al.
O
TP63
KRT5
Counts
EN1 low
EN1 high
Peng et al.

**Figure S2**

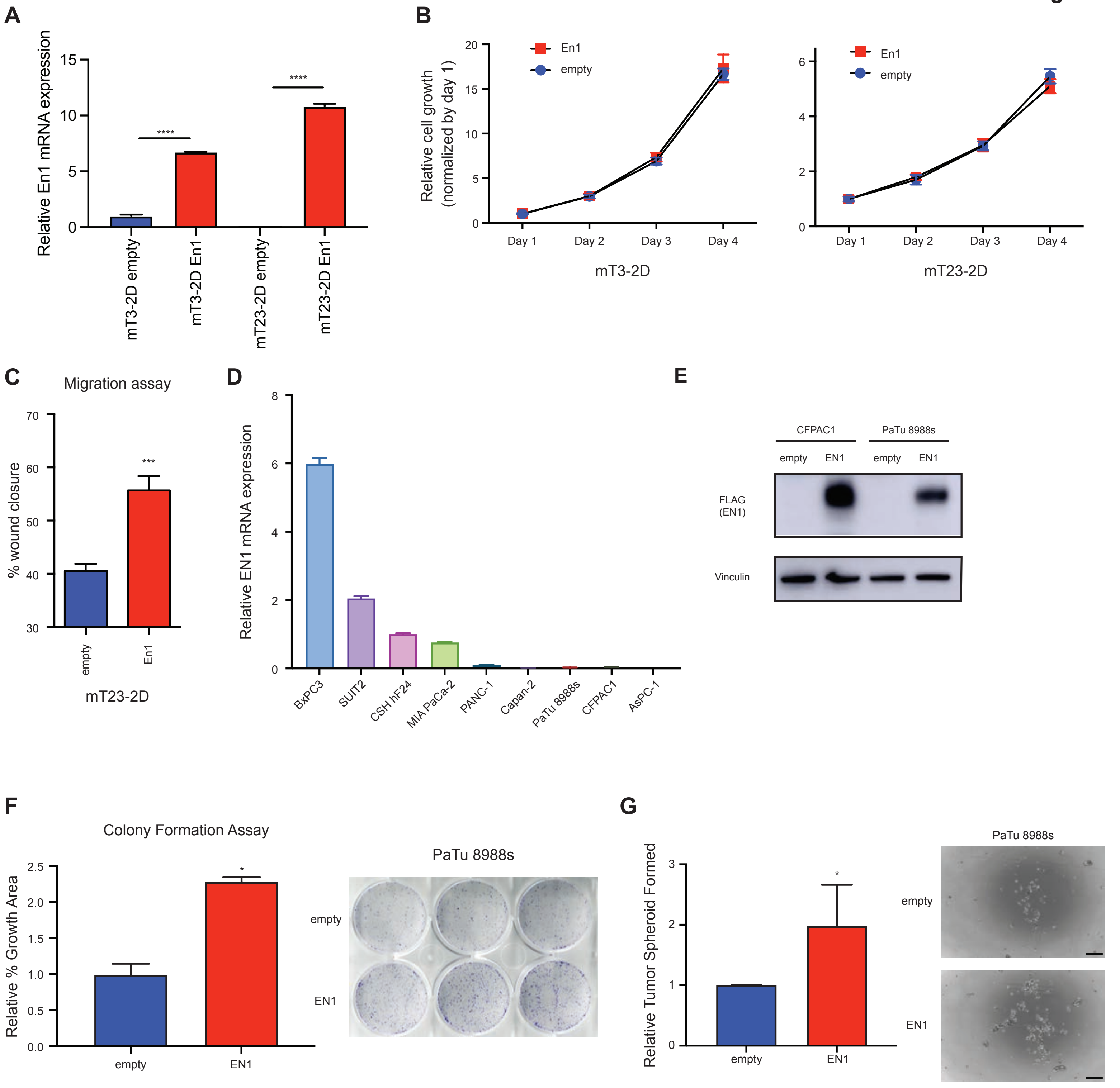
A
Relative En1 mRNA expression
****
****
mT3-2D empty
mT3-2D En1
mT23-2D empty
mT23-2D En1
B
Relative cell growth
(normalized by day 1)
En1
empty
Day 1
Day 2
Day 3
Day 4
mT3-2D
mT23-2D
C
Migration assay
% wound closure
***
empty
En1
mT23-2D
D
Relative EN1 mRNA expression
BxPC3
SUIT2
CSH hF24
MIA PaCa-2
PANC-1
Capan-2
PaTu 8988s
CFPAC1
AsPC-1
E
CFPAC1
PaTu 8988s
empty
EN1
FLAG
(EN1)
Vinculin
F
Colony Formation Assay
Relative % Growth Area
PaTu 8988s
empty
EN1
G
Relative Tumor Spheroid Formed
PaTu 8988s
empty
EN1

### Figure S3

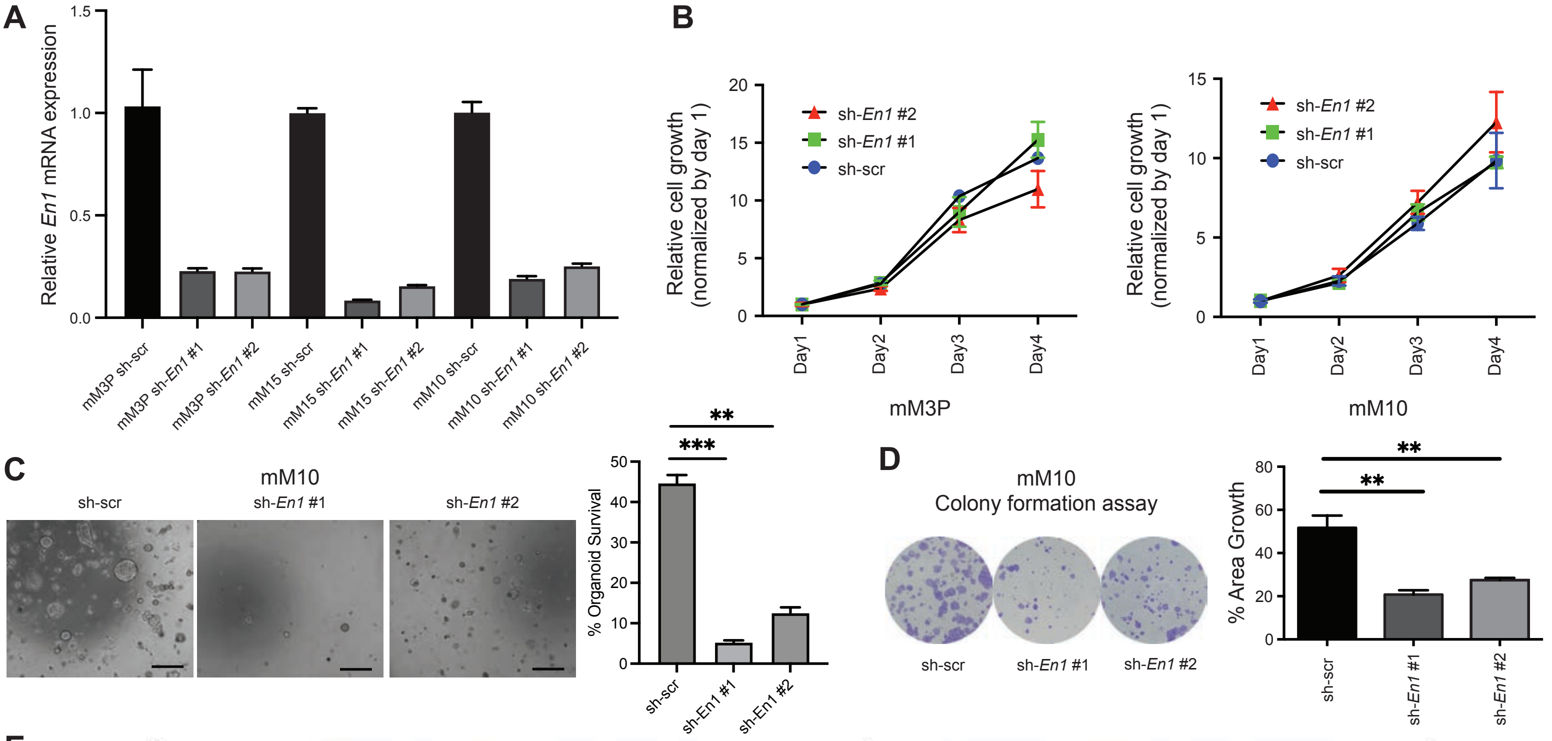


E

| Case | Metastases from M3P/sh-scr orthotopic transplants #1 | #2 | #3 | #4 | #5 | Metastases from M3P/shEn1 orthotopic transplants #1 | #2 | #3 | #4 |
|---|---|---|---|---|---|---|---|---|---|
| Liver | Met | Met | Met | Met | Met | No | No | Met | No |
| Lung | Micromets (>10) | Micromets (>10) | Micromets (>10) | Micromets (>10) | Micromets (<10) | Micromets (<10) | Micromets (>10) | Micromets (<10) | Micromets (<10) |
| Diaphragm | Met | Met | Met | Met | No | No | Met | Met | No |
| Ovary | Met | No | No | No | No | No | No | No | No |
| Mesentery | Met | Met | Met | No | No | No | Met | Met | No |
| Stomach | Met | No | No | No | No | No | No | No | No |
| Peritoneum | No | No | No | No | Met | No | Met | Met | No |

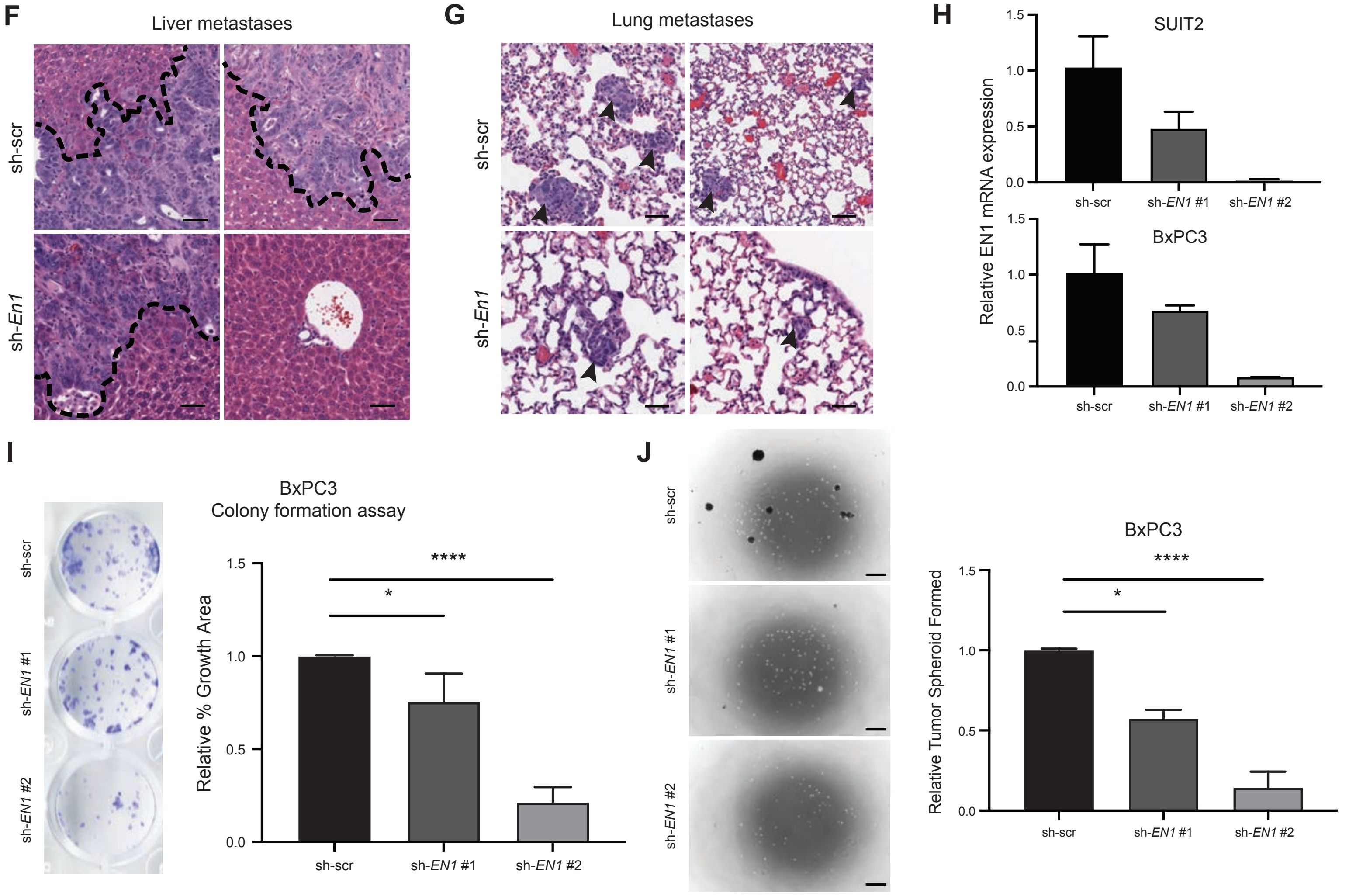

#### Figure S4

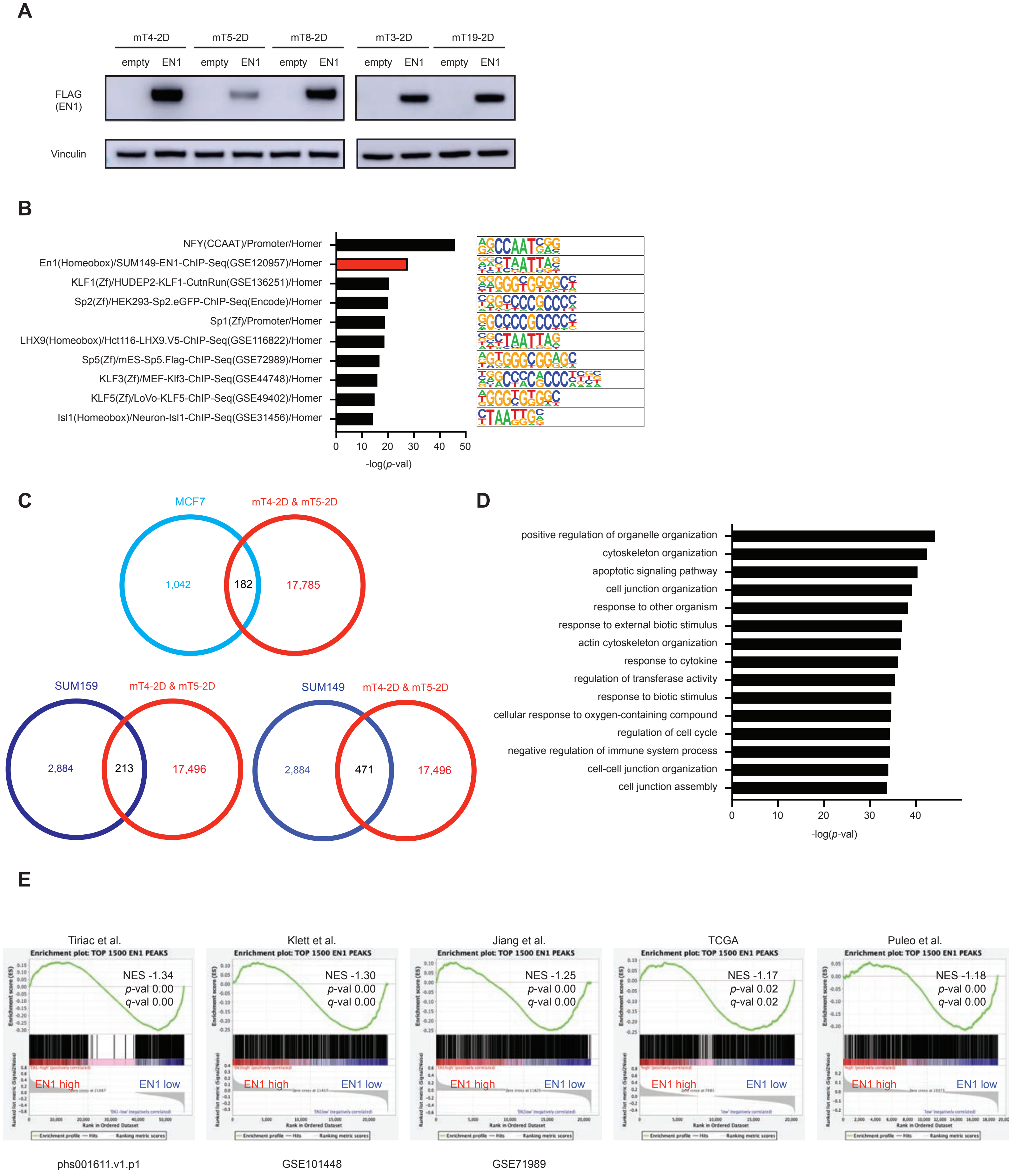
A
mT4-2D
mT5-2D
mT8-2D
mT3-2D
mT19-2D
empty EN1 empty EN1 empty EN1 empty EN1 empty EN1
FLAG (EN1)
Vinculin
B
NFY(CCAAT)/Promoter/Homer
En1(Homeobox)/SUM149-EN1-ChIP-Seq(GSE120957)/Homer
KLF1(Zf)/HUDEP2-KLF1-CutnRun(GSE136251)/Homer
Sp2(Zf)/HEK293-Sp2.eGFP-ChIP-Seq(Encode)/Homer
Sp1(Zf)/Promoter/Homer
LHX9(Homeobox)/Hct116-LHX9.V5-ChIP-Seq(GSE116822)/Homer
Sp5(Zf)/mES-Sp5.Flag-ChIP-Seq(GSE72989)/Homer
KLF3(Zf)/MEF-Klf3-ChIP-Seq(GSE44748)/Homer
KLF5(Zf)/LoVo-KLF5-ChIP-Seq(GSE49402)/Homer
Isl1(Homeobox)/Neuron-Isl1-ChIP-Seq(GSE31456)/Homer
0 10 20 30 40 50
-log(p-val)
C
MCF7
mT4-2D & mT5-2D
1,042
182
17,785
SUM159
mT4-2D & mT5-2D
2,884
213
17,496
SUM149
mT4-2D & mT5-2D
2,884
471
17,496
D
positive regulation of organelle organization
cytoskeleton organization
apoptotic signaling pathway
cell junction organization
response to other organism
response to external biotic stimulus
actin cytoskeleton organization
response to cytokine
regulation of transferase activity
response to biotic stimulus
cellular response to oxygen-containing compound
regulation of cell cycle
negative regulation of immune system process
cell-cell junction organization
cell junction assembly
0 10 20 30 40
-log(p-val)
E
Tiriac et al.
Enrichment plot: TOP 1500 EN1 PEAKS
NES -1.34
p-val 0.00
q-val 0.00
EN1 high
EN1 low
Klett et al.
Enrichment plot: TOP 1500 EN1 PEAKS
NES -1.30
p-val 0.00
q-val 0.00
EN1 high
EN1 low
Jiang et al.
Enrichment plot: TOP 1500 EN1 PEAKS
NES -1.25
p-val 0.00
q-val 0.00
EN1 high
EN1 low
TCGA
Enrichment plot: TOP 1500 EN1 PEAKS
NES -1.17
p-val 0.02
q-val 0.02
EN1 high
EN1 low
Puleo et al.
Enrichment plot: TOP 1500 EN1 PEAKS
NES -1.18
p-val 0.00
q-val 0.00
EN1 high
EN1 low
phs001611.v1.p1
GSE101448
GSE71989

### Figure S5

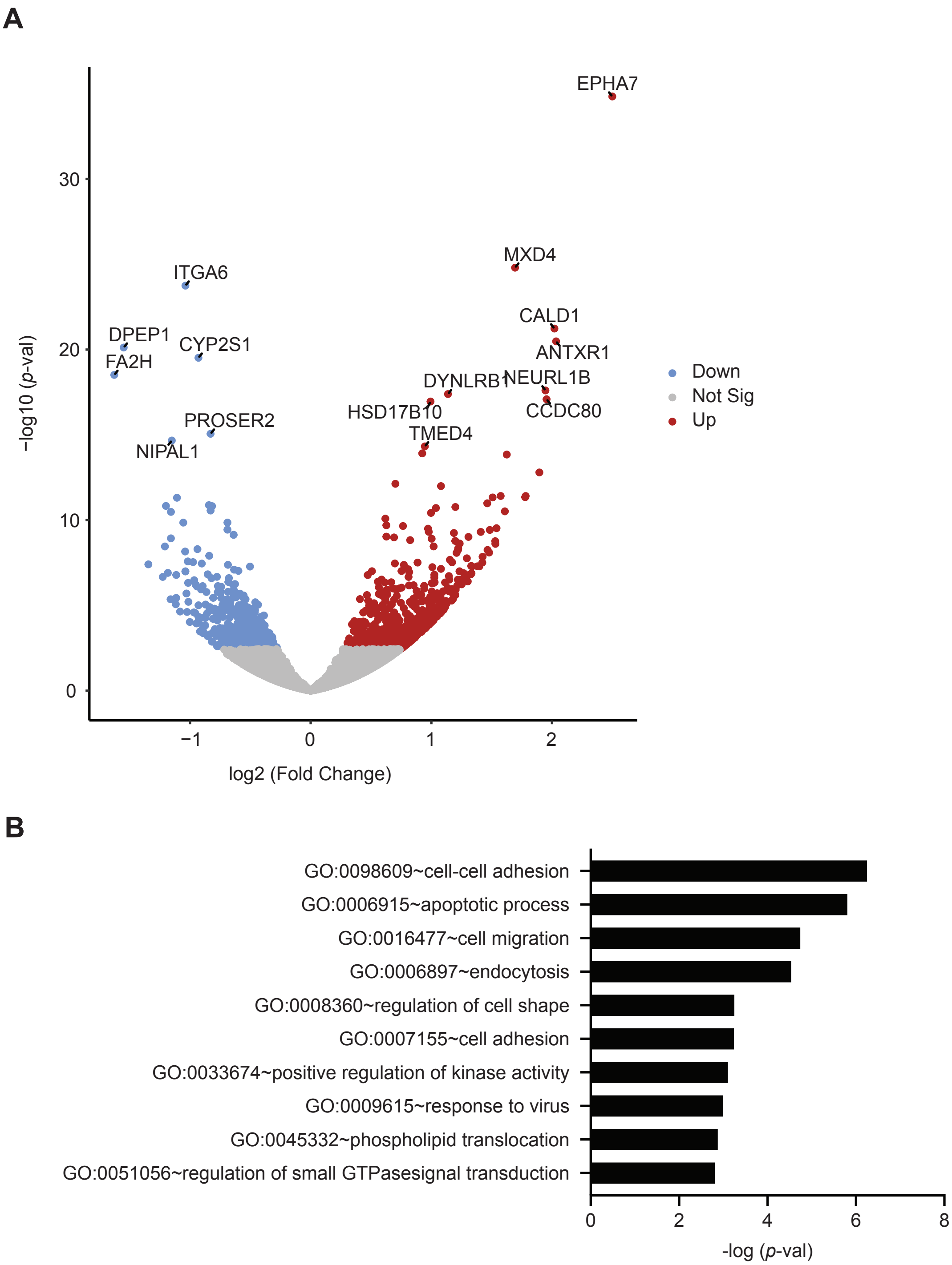
A
EPHA7
MXD4
ITGA6
CALD1
DPEP1
CYP2S1
ANTXR1
FA2H
DYNLRB1
NEURL1B
HSD17B10
CCDC80
PROSER2
TMED4
NIPAL1
Down
Not Sig
Up
−log10 (p-val)
log2 (Fold Change)
30
20
10
0
−1
0
1
2
B
GO:0098609~cell-cell adhesion
GO:0006915~apoptotic process
GO:0016477~cell migration
GO:0006897~endocytosis
GO:0008360~regulation of cell shape
GO:0007155~cell adhesion
GO:0033674~positive regulation of kinase activity
GO:0009615~response to virus
GO:0045332~phospholipid translocation
GO:0051056~regulation of small GTPasesignal transduction
0
2
4
6
8
-log (p-val)

### Figure S6

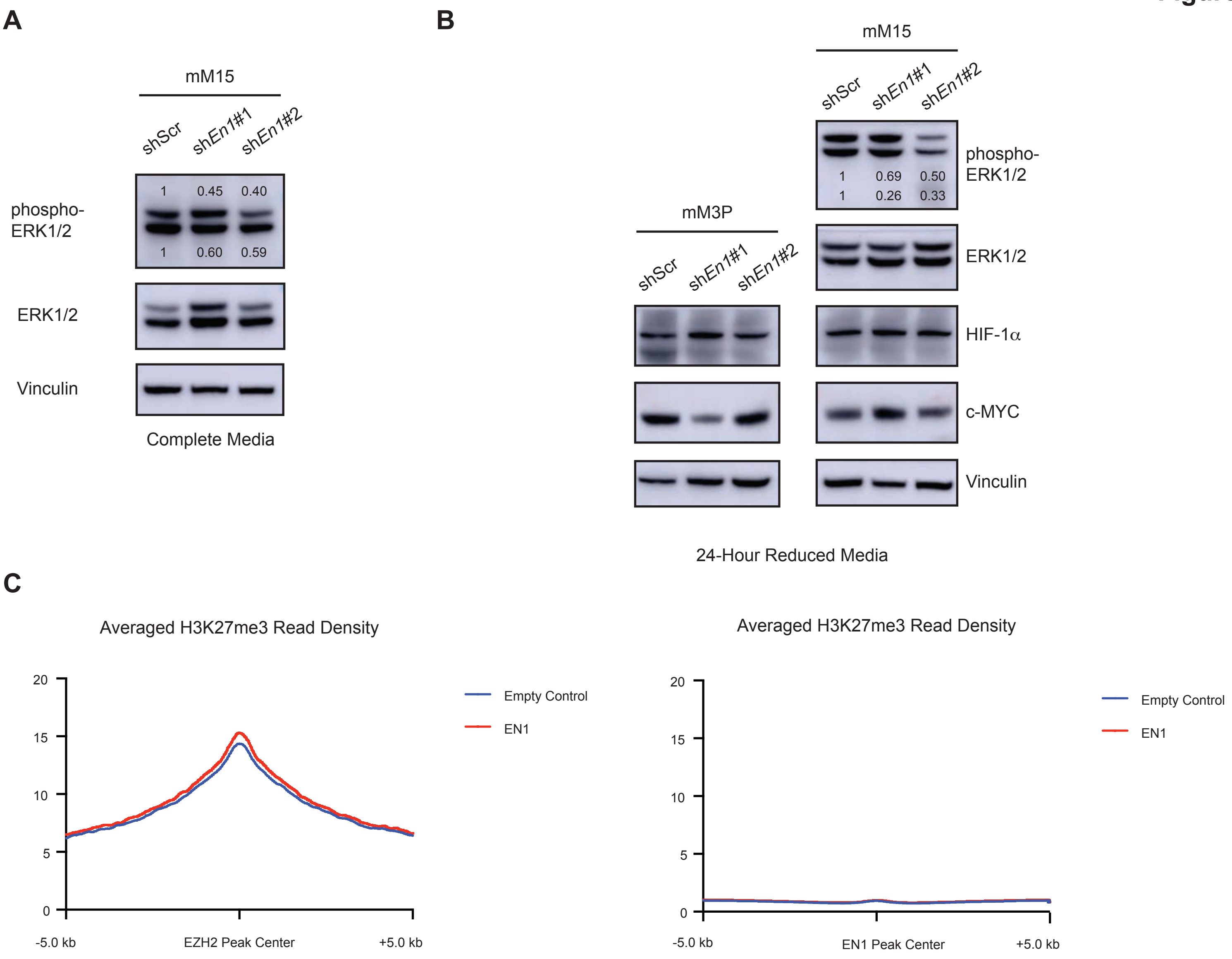
A
mM15
shScr
shEn1#1
shEn1#2
phospho-ERK1/2
1 0.45 0.40
1 0.60 0.59
ERK1/2
Vinculin
Complete Media
B
mM3P
shScr
shEn1#1
shEn1#2
mM15
shScr
shEn1#1
shEn1#2
phospho-ERK1/2
1 0.69 0.50
1 0.26 0.33
ERK1/2
HIF-1α
c-MYC
Vinculin
24-Hour Reduced Media
C
Averaged H3K27me3 Read Density
20
15
10
5
0
-5.0 kb
EZH2 Peak Center
+5.0 kb
Empty Control
EN1
Averaged H3K27me3 Read Density
20
15
10
5
0
-5.0 kb
EN1 Peak Center
+5.0 kb
Empty Control
EN1

**Figure S7**

**A**

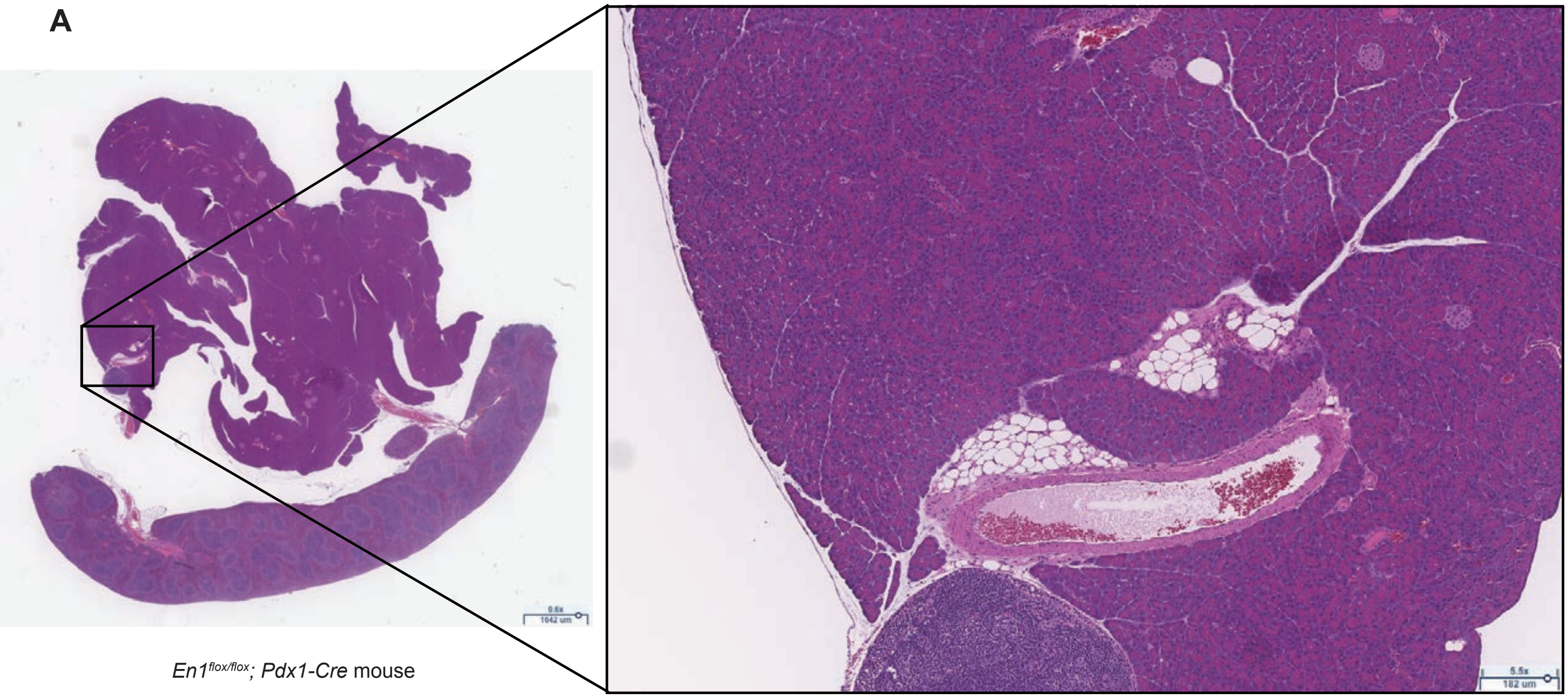

0.6x
1642 um
5.5x
182 um
*En1*$^{flox/flox}$*; Pdx1-Cre* mouse


**B**

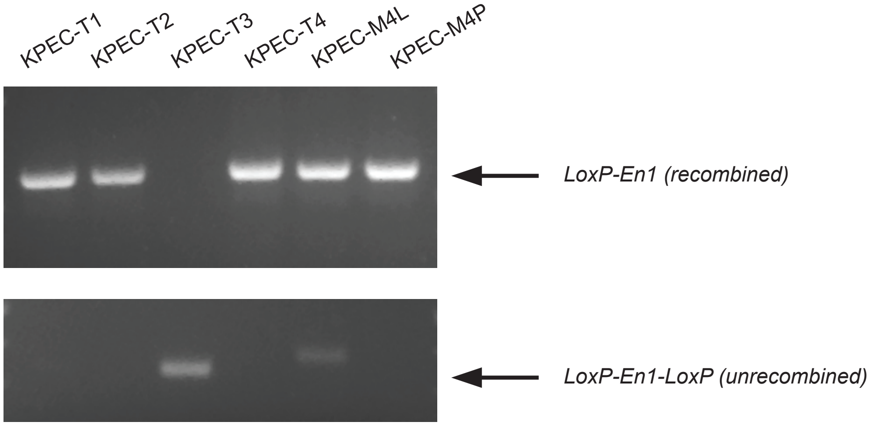

KPEC-T1
KPEC-T2
KPEC-T3
KPEC-T4
KPEC-M4L
KPEC-M4P
*LoxP-En1 (recombined)*
*LoxP-En1-LoxP (unrecombined)*
